# Structural basis for hydrolytic splicing of a circularly permuted group II intron

**DOI:** 10.64898/2025.12.30.696962

**Authors:** Xiaobin Ling, Yuqi Yao, Dmitrij Golovenko, Kangkang Song, Chen Xu, Jianhua Gan, Zhenguo Chen, Jinbiao Ma

**Author notes:** These authors contributed equally (Xiaobin Ling, Yuqi Yao, Dmitrij Golovenko). Correspondence should be addressed to X.L., Z.C., J.M.

## Abstract

Group II introns are self-splicing ribozymes that are considered the ancestors of the eukaryotic spliceosome. Unlike canonical group II introns that self-splice to generate linear exons and lariats, circularly permuted (CP) group II introns identified in various bacterial phyla perform back-splicing, resulting in the production of circular RNAs and branched products via branching pathway. Furthermore, CP introns may switch to a hydrolysis pathway resulting in distinct products to differentially regulate retrotransposition. In this study, we present biochemical data and high-resolution cryogenic electron microscopy (cryo-EM) structures of a CP group II intron from *Comamonas testosteroni* KF-1 (*Cte* 1), allowing mechanistic dissection of the switch from the branching pathway to the hydrolysis pathway, and enabling reconstruction of both steps of the hydrolysis pathway. The structures reveal that CP group II intron undergoes the hydrolysis pathway upon mutations of the branch point or splice sites (SS) due to rearrangements in the active site. Here, the branching nucleotide in domain D6 is retracted from its catalytically competent conformation, giving way to the nucleophilic water molecule to attack the 5′ splice site. Furthermore, we visualized the intermediates of the second splicing step, which reveal the movement of domain D6 out of the way for the 3′-splicing to occur, closely resembling the second-step structures of the branching pathway. Finally, our structures provide direct evidence for domains D1–D3 acting as a scaffold in group II introns. Together, these findings visualize the complete hydrolysis pathway and offer a new strategy to engineer CP group II introns for circular RNA production, with potential applications in both basic research and therapeutic development.

## Introduction

Group II introns are mobile ribozymes that can self-splice from RNA precursors and reverse splice into DNA, thereby propagating themselves in bacterial and organellar genomes^1,2^. They are considered the ancestors of spliceosome introns and retrotransposons in eukaryotes, thus providing a model for understanding the mechanism and regulation of these related processes^3^. A typical splicing reaction of group II introns and spliceosomes involves two sequential transesterification reactions. The initial step involves a nucleophilic attack by the 2′-OH group of an internal adenosine (the “branch point”) at the 5′ splice junction, resulting in the formation of a 2′-5′ phosphodiester branch within the intron. In the second step, the 3′-OH of the cleaved 5′ exon attacks the 3′ splice site, thus joining the two exons with a 3′-5′ phosphodiester bond and releasing the intron as a lariat. For most group II introns, the first step is rate-limiting^4,5^.

An important alternative to the “branching” pathway of splicing is the hydrolytic splicing, where a water molecule acts as a nucleophile instead of the 2′-OH of the branch point adenosine, resulting in lariat-free products ^6,7^. Hydrolytic splicing occurs in vivo^7,8^, and was characterized *in vitro* using constructs with mutations at or around the branch point^9,10^. The branching and hydrolytic pathways were proposed to underlie the regulation of group II intron activity. First, because hydrolytic splicing is irreversible, its linear intron product cannot reverse-splice into DNA for further genomic propagation. Second, the linear intron may allow translation of intron-encoded proteins (IEPs), facilitating subsequent splicing, reverse-splicing^5^, and/or other processes.

Group II introns are categorized into three main classes—IIA, IIB, and IIC—based on the sequence of their IEPs and RNA secondary structural features. Group IIA and IIB introns readily form branched lariats, whereas group IIC introns predominantly splice via the hydrolysis pathway^2^. Despite their differences, they all share a conserved arrangement of six structural domains (D1–6). The largest domain D1 enables exon binding sites (EBS) recognition by base pairing with intron binding sites (IBS) on the exon. D2 and D3 provide additional structural stability through interactions with D5 and D6 and notably, a conserved intervening junction (J2/3) that is part of the catalytic core. D4 often encodes an IEP that facilitates self-splicing and reverse-splicing. D5 docks into D1 and contains the most conserved nucleotides that form the active sites. D6 provides the branch point adenosine for the first transesterification step^2,6^.

Substantial progress in the structural understanding of group II intron splicing and reverse-splicing revealed a conserved catalytic center among different classes of group II introns, which is used in both steps of splicing^11–20^. The active site for the pre-first step of splicing is almost identical between the branching and hydrolysis pathway^11,19^, raising the question of how the choice between the two pathways is made. Several artificial and natural mutations have been reported to convert the branching pathway to the hydrolysis pathway^7–9^, but a structural basis for this conversion is lacking. Interestingly, when the D1–5 of the hydrolytic group IIC intron from *O. i* is fused to D6 from *A.v.* group II intron, the branching pathway is activated, suggesting that D6 provides the critical structural features determining the choice of the pathway^21^. While recent cryo-EM studies have provided many details of splicing through the branching pathway, structural analysis of the hydrolysis pathway is limited to early crystallographic studies, in which D6 conformations were not resolved^11,12^.

A family of naturally occurring, circularly permuted (CP) group II intron was recently identified in diverse bacterial genomes. These CP group II introns contain rearranged structural domains and produce circular RNAs (circRNAs) via back-splicing^22^. We have reported high-resolution cryo-EM structures of the 3′ truncated CP group II intron from *Comamonas testosteroni* KF-1 (*Cte* 1), showing that splicing of this intron shares the conserved catalytic mechanism with group II introns through the branching pathway, facilitated by an auxiliary sequence (AUX). Here, we determined structures of *Cte* 1 mutants, which utilize the hydrolysis pathway, capturing the steps of this pathway. Our work reveals how the intron employs unconventional D6 docking into D1 to route the splicing via the hydrolysis pathway. D1–3 domains rearrange to accomplish the second step of the hydrolysis pathway. Finally, the two hydrolysis products reveal the structural rearrangements leading to the circular RNA produced.

## Results

### Conversion from the branching to hydrolysis pathway in *Cte* 1 mutants

To compare the CP introns undergoing the branching and hydrolytic pathways, we first determined the full-length wild-type structure of group II intron from *Cte 1* undergoing the branching (Fig. 1a, 1b). Cryo-EM classification revealed the conformation representing the steps of pre-splicing (Pre, 2.75 Å; Fig. 1c and Extended Data Fig. 2). The overall conformations are similar to that in our previous cryo-EM structures of *Cte* 1, which lacked the D4 domain^23^. D4 is thought to interact with the ORF encoding the IEP and to facilitate the interaction between IEP and the ribozyme^17–19^. Stabilized by D4, the neighboring subdomains P1 and P2 dock at D2 and D3 via long range interactions, as predicted previously^22^, stabilizing the overall conformation of the Pre state (Fig. 1c and Extended Data 6). Furthermore, G809–C813 of D4 base-pair with C50–C54 of D5 (Extended Data Fig. 6a). Our structures therefore reveal that D4 contributes to the stabilization of CP group II intron in the absence of IEP.

**Figure 1.**
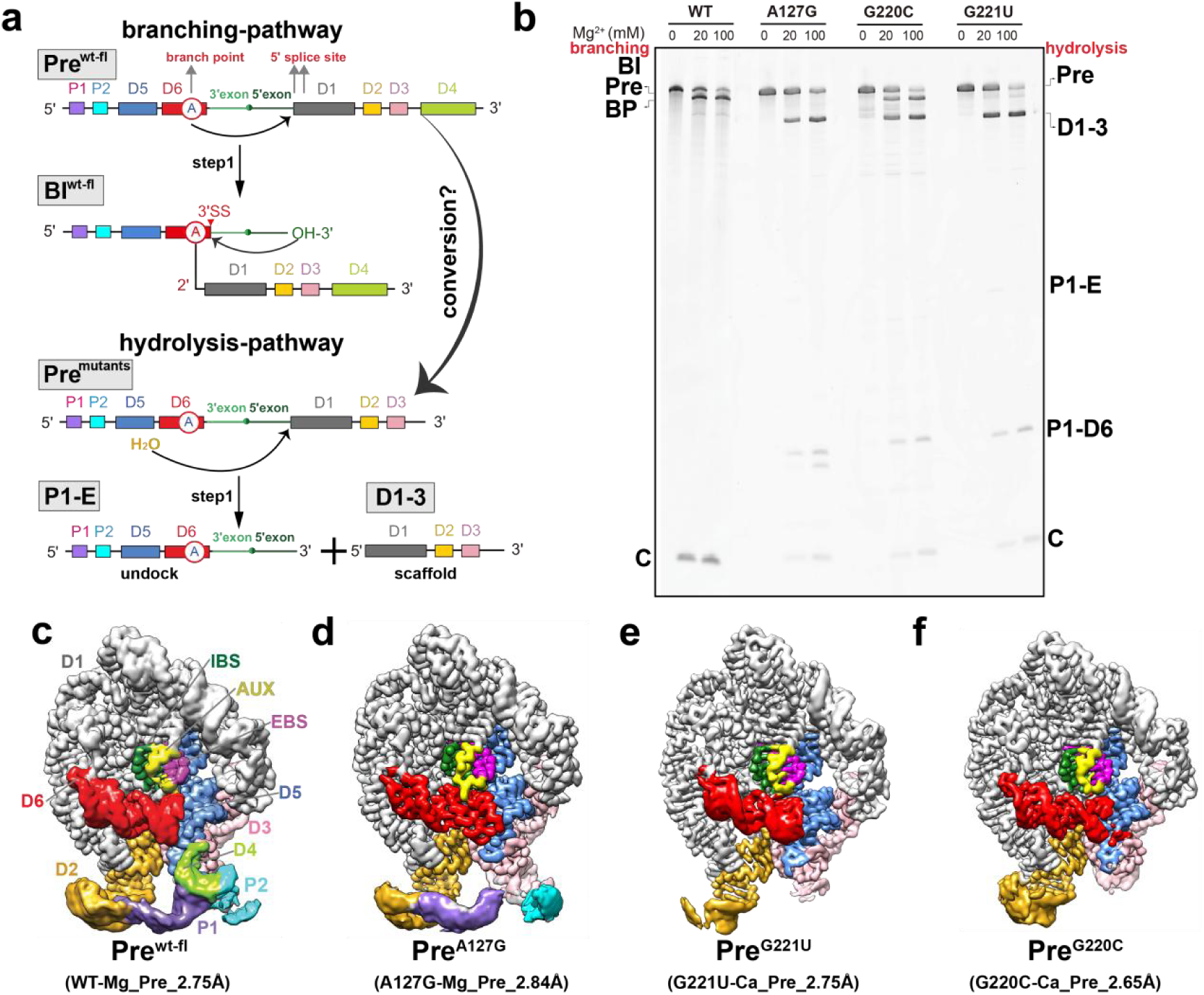
Cryo-EM structures of hydrolytic *Cte*mutants. **a.** Cartoon of the 1^st^ step of branch-pathway and hydrolysis-pathway of *Cte* group II intron back-splicing. **b.** The Syber Gold-stained 6% urea–polyacrylamide gel electrophoresis (PAGE) gels showing different mutants back-splicing activity by branch pathway or hydrolysis pathway at 20 or 100 mM MgCl_2_ concentration. WT and A127G samples used to cryo-EM. **c-f.** The Pre states of cryo-EM map and molecular model of the *Cte* 1_fl and branch point mutant A127G, 5*’* SS mutant G220C and G221U.

To address the mechanism of the hydrolysis pathway, we identified *Cte* 1 mutants that partially or completely employ the hydrolysis pathway for the first step of splicing. The branch-point mutation A127G and the near-5′-splice site (5′-SS) mutant G221U switch the enzyme to the hydrolysis pathway without producing notable branching intermediates, whereas the 5′-SS mutant G220C yields a mixture of the hydrolysis and branching products (Fig. 1b and Extended Data Fig. 1a–c).

We next determined cryo-EM structures of these constructs at 2.6–2.8 Å resolution, sufficient to visualize the structural details underlying active-site differences (Fig. 1d–f and Extended data Fig. 3–5). To capture the early Pre-splicing states, we determined structures of G220C and G221U mutants in the presence of CaCl_2_ to inhibit the splicing activity (Fig. 1e, 1f, Extended Data Fig. 1d, 4 and 5). We were also able to capture the first step of hydrolysis in the Pre state using the A127G mutant in the presence of MgCl_2_ (Fig. 1b and Extended data Fig. 3). In contrast to the WT construct, which produced the BI intermediate under the same conditions including MgCl_2_ (Fig. xx), the A127G and other mutants therefore bring mechanistic insights into the hydrolysis pathway.

### Active-center rearrangements upon D6 conformational change lead to the hydrolysis pathway

For all structural alignment we used D5 as an anchor. Notably, there is a significant shift in the D6 conformation (Extended Data Fig. 7b). The A127G mutation resulted in a displacement of 7.8 Å, whereas the G221U mutation causes a displacement of approximately 3.8 Å. The displacement difference between A127G and G221U is 6.5 Å. In all three mutants, the active sites adopt different conformations, but they have two common features (movie S1 and movie S2). First, the G220 loop retains its sharply kinked conformation with phosphate 220 at the tip coordinating either one or two cations, resembling the conformation of the pre-attack position in the WT structure (Fig. 2a). Second, the branching nucleotides 127, guanosine in the A127G mutant or A127 in the other mutants, are found in different positions away from G220 (Fig. 2). Unlike in the WT Pre structure, where the 2′-OH 127 is placed 3.3 Å from phosphorus 220, the respective distances range from 4 Å in the G220C mutant to 7.5 Å in G221U and to ∼12.5 Å in A127G. In these conformations, the G220 loop remains an attractive nucleophilic target, whereas the nucleophilic 2′-OH group is detached, providing space for a smaller nucleophile—such as water—to attack the G220 phosphate group. Indeed, the separation of 2′-OH in WT Pre and our mutant structures correlates with the extent of the hydrolysis pathway, as the most closely positioned hydroxyl in WT undergoes the branching pathway, the slightly shifted G220C demonstrates both the branching and hydrolysis intermediates, while the remaining most separated mutants undergo the hydrolysis pathway (Fig. 1b and Fig. 2).

**Figure 2:**
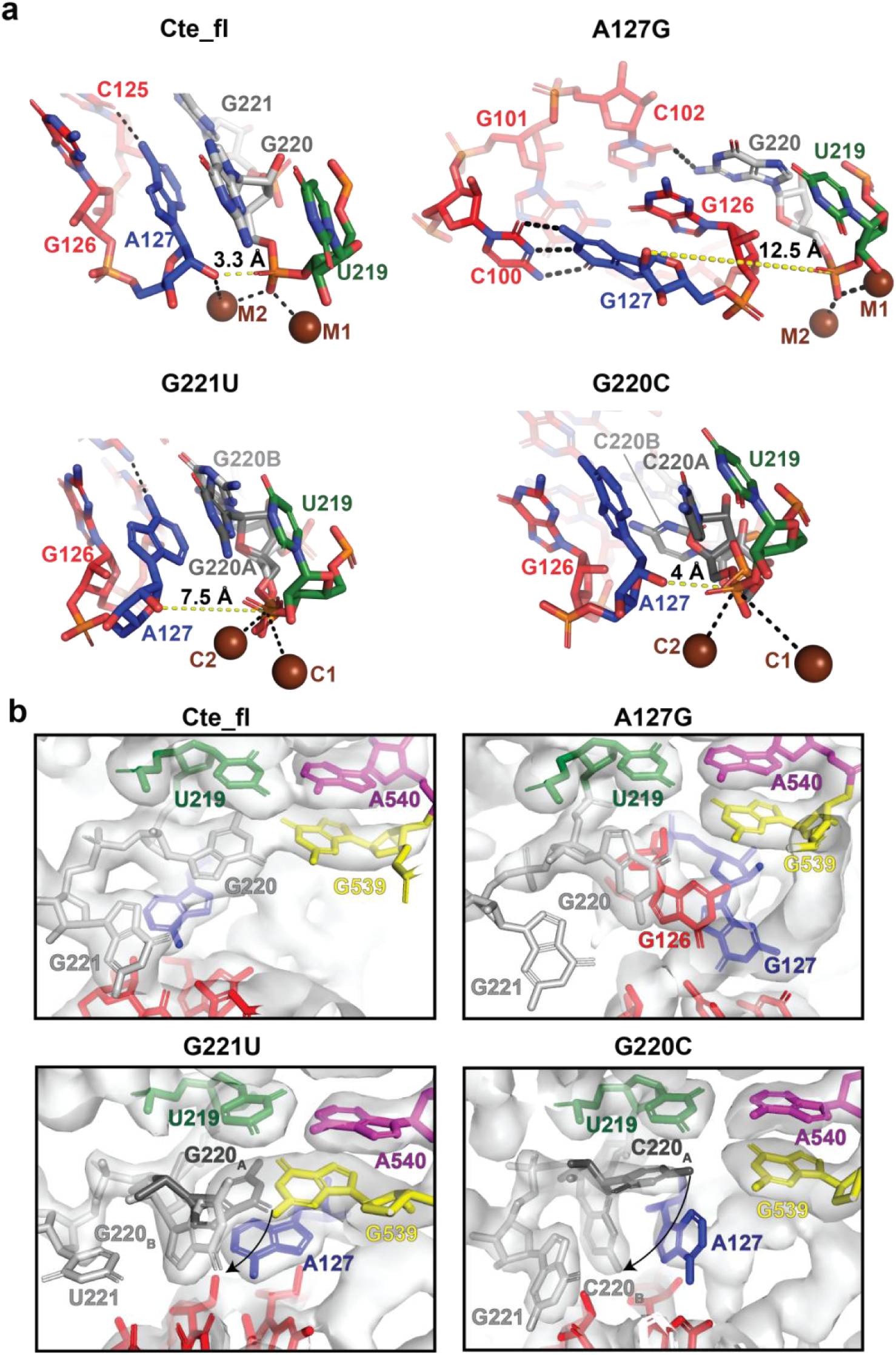
The catalytic center of hydrolysis pathway mutants in Pre states. **a–b.** The catalytic center of *Cte* full length (Cte-fl), mutants A127G, G221U and G220C, showing the converstion of the branch pathway to the hydrolysis pathway.

In all mutants, the rearrangement of the active site occurs due to the differences in the nucleotide identities next to the branch point that disrupt the interactions present in the WT Pre structures (movie S1 and movie S2). In the G220C mutant, C220 is unstacked from A127, consistent with the less favorable stacking of pyrimidines on purines than purines on purines^24^. This results in the ∼2 Å shift of the A127 ribose, placing its hydroxyl group farther from the scissile phosphate of G220 than in the WT Pre structure (Fig. 2a). In the G221U mutant, U221 is also separated from A127, in contrast to WT Pre, where G221 participates in a triple-pairing with the D6 helix and stacks on A127. Here, A127 is shifted further from its position in WT Pre, placing its ribose ∼7 Å away from the G220 phosphate (Fig. 2a). In A127G, the mutated residue G127 acquires a new base-pairing capability, which brings it into the D6 helix to Watson-Crick pair with C100 (Fig. 2a). This base pair is further stabilized by being sandwiched between Watson-Crick pairs G126-C538 and G99-C128. Due to the large rearrangement of the branching position, the D6 helix in the A127G mutant is less well ordered than in other structures, indicating D6 mobility as described in Supplementary Discussion.

### Changes in δ-δ′ interaction in the conversion from branching to hydrolysis pathway

Through density analysis, we captured the δ-δ′ interaction between G220 and G539 in both the G220C and G221U mutants, which is crucial for correctly recognizing the 5′-SS in the branching pathway. During the conformational shift from branching to hydrolysis, however, notable differences were observed (movie S1). 220 of G220C and G221U mutants have two conformations which are named G220_A_ and G220_B_. In the G221U mutant, G220_A_ does not exhibit strong residual branching density associated with G539; instead, G220_B_ dissociates from G539 and inserts into the minor groove of D6. In contrast, in the G220C mutant, two strong density signals are present. In the first, C220_A_ base-pairs with G539 during branching, while the other, C220_B_, points downward, moving away from G539 and creating sufficient space for the branch point A127 to adopt a new conformation. In the A127G mutant, G220 stabilizes its conformation by stacking with A127 and G126, thus distancing itself from G539 (Fig. 2b). Overall, in all three mutants, G220 or C220 disrupts the δ-δ′ interaction between G220 and G539 that is formed during branching and instead establishes new structural stabilizing interactions. These changes facilitate the transition from the branching to the hydrolysis pathway.

### Conformational changes of the triple helix and metal cluster

The triple helix provides a stable platform for organizing the hydrolytic active site. Upstream of the 5*’*-SS, stability is mediated by base pairing with the EBS; downstream, metal cluster coordination—particularly at K2—supports the conformation necessary for nucleophilic attack^12^. Using the *O. iheyensis* group II intron structure (PDB 4FAQ) as a reference, we defined K2 as the density between the phosphate groups of G224 and A695 (Extended Data Fig. 8). In the wild-type, no extra downstream density is observed. In contrast, all three hydrolysis-prone mutants exhibit additional density near the 5′-SS, though distance measurements exclude assignment to K^+^ or Mg^2+^ (data not shown).

The wild-type and 4FAQ structures are highly similar, with the main difference being a downward shift of K2 to G224, corresponding to G5 in 4FAQ. In the wild-type, K2 stabilizes the 5*’*-SS terminal base through interaction with A695 rather than the 2-nt bulge. Apart from this, the architecture remains nearly identical (Extended Data Fig. 8a). Comparison to 4FAQ highlights a change in domain D6. In the wild-type, G221 engages with D6 via π-π*’* interactions, akin to U2 in 4FAQ, reducing the reliance on K2 for stabilization. The absence of D6 in 4FAQ shifts K2 upward to stabilize multiple downstream bases, introducing a conformational tilt.

We next analyzed the triple helix and metal cluster rearrangements in the mutants (movie S2). For A127G mutant, G221 adopts two conformations: an outward-flipped (branching-like) state (G221_A_) and an inward state stacking with A695 and U82 (G221_B_). K2 density is weak or absent, and stacking likely compensates for the loss of metal-mediated stabilization. For G221U mutant, J2/3 distortion disrupts stacking among A697–G696, G83, and C289–G288/C377, destabilizing the branching configuration and favoring hydrolysis. For G220C mutant, the triple helix and metal cluster rearrangements of both the wild-type and 4FAQ are preserved, maintaining competence for both pathways (Extended Data Fig. 8b–d).

### Visualizing the second step of the hydrolysis pathway *in trans*

Due to the unique inverted structure of the D5–D6 domain in the CP group II intron, the first step of the hydrolytic pathway generates two linear RNA products: D1-D2-D3 (D1–3) and P1-P2-D5-D6-Exon (P1–E) (Fig. 1a)^22^. Our cryo-EM analysis of the A127G mutant incubated with MgCl_2_ yielded a 3. 1Å map corresponding to the D1–3 structure (Fig. 1b and 3c, Extended Data Fig. 3, movie S3). The conformation of these domains is very similar to those in the pre-reaction full-length intron, consistent with the idea that the large D1 and smaller D2–3 domains form a scaffold for the splicing reaction^2^. The second step of the hydrolysis pathway was proposed to also depend on the D1 scaffolding^2^, likely required to organize the splice site for freeing the exon from the P1–D6 domains. Because P1–E (product of the first step) and P1–D6 (product of the second step) are much smaller than the D1-containig molecules, we did not identify them in the cryo-EM dataset. Our biochemical assays suggest that the conversion of P1-E to P1-D6 in the A127G mutant is fast and coincident with the formation of D1–3 (Fig. 1b), consistent with previous studies^2,12^. It is therefore likely that D1–3 in the A127 mutant dataset arose as the result of complete hydrolytic splicing and subsequent intron disassembly, consistent with the requirement of D1–3 for both steps (Fig. 1b, 3c, 3d).

**Figure 3:**
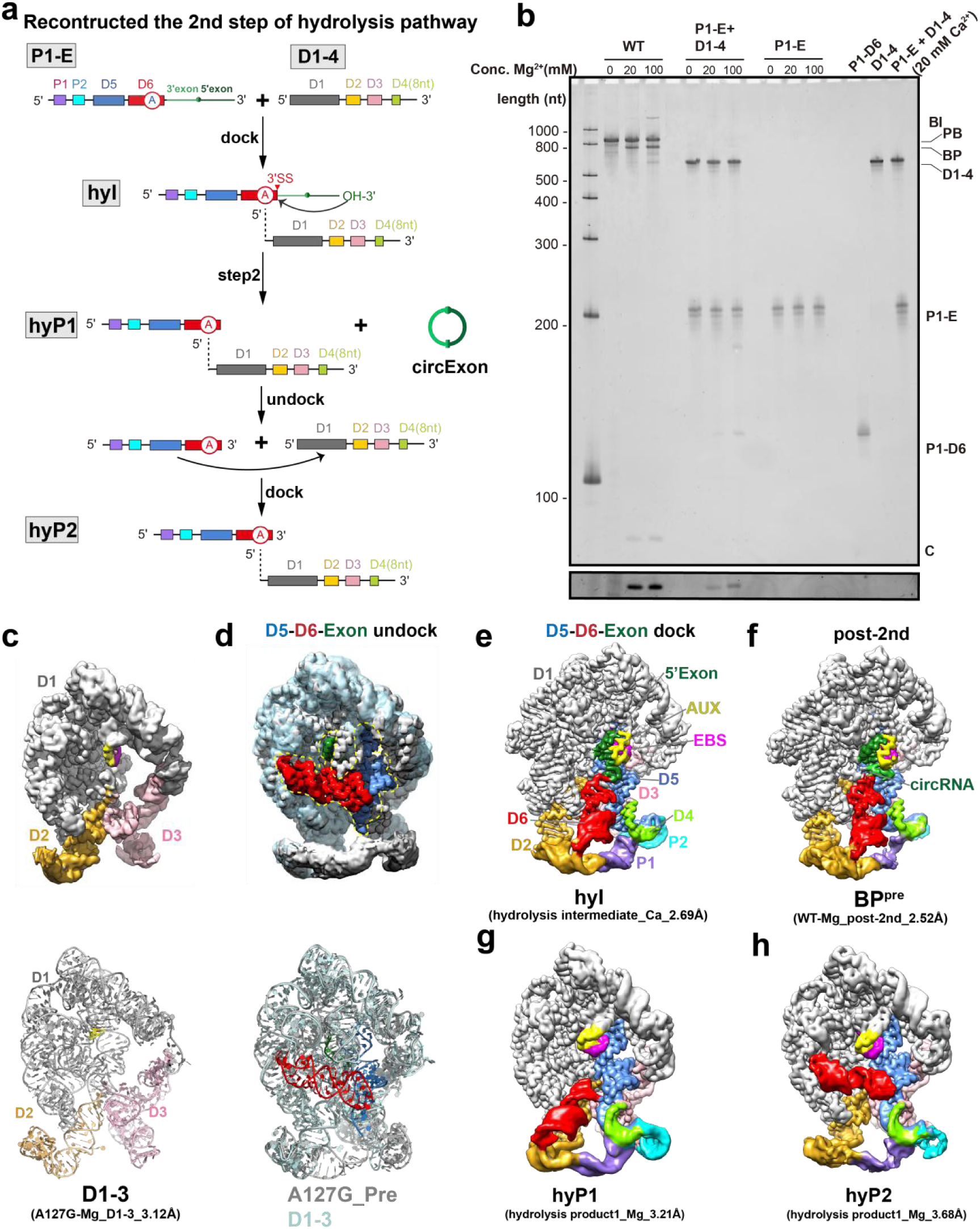
Reconstructed 2^nd^ step of hydrolysis via “trans-splicing”. **a.** The schematic of the 2^nd^ step of CP group II intron to produce circular RNA by “trans-splicing”. **b.** Biochemical analysis of trans-splicing between P1-E and D1-4. The Syber Gold-stained 6% urea–polyacrylamide gel electrophoresis (PAGE) gels shows that P1-E docked with D1-4 to produce circular exon. The same reaction was used in cryo-EM sample preparation. **c.** The cryo-EM map and model of D1-3. **d.** The aligned cryo-EM maps and models shows that the D5-D6-Exon may undocked from 1^st^ step structure. **e-f.** Cryo-EM maps of the *Cte* 1 hydrolysis intermediate (hyI) and the pre-of branching product (BP^pre^) states. g-h. The cryo-EM maps of hydrolysis product 1 and 2.

To visualize the fast second step of splicing, we combined artificially transcribed products of the first step of the hydrolysis reaction, P1–E and D1–4, in the presence of inhibitory CaCl_2_ to prevent the fast conversion to final products (movie S2). In the hydrolysis intermediate (hyI) structure corresponding to the 2^nd^ step of the hydrolysis pathway, the 3′–5′ phosphodiester bond has formed by U219 and C135, the circular exon was retained in the structure (Fig. 3e, 4a, Extended data Fig. 9). This conformation confirmed the process of P1-Exon truly could docking back into D1–4 structure.

Compared with the pre-branching product (BP^pre^) conformation, where a covalent 3′-5′ phosphodiester bond has formed (Fig. 3f), the catalytic active sites of BP^pre^ and hyI are generally similar (Fig. 4a). We previously showed that AUX is a Auxiliary sequence forming triple helix with IBS-EBS in Cte 1, which function is similar with the IEP of group II intron. To investigate the roles of C538 and G539 in AUX during the transition from the 1^st^ step to the 2^nd^ step of the hydrolysis pathway, we compared the conformational changes between the WT hyI state and the Pre states of the three mutants (Fig. 4b). In the branching pathway, C538 undergoes a ∼90° downward flip from the Pre to the BP^pre^ state, and the G–G interaction between G539 and G220 is replaced by a Watson-Crick base-pair between G539 and C134. In the hydrolysis pathway, except for the A127G mutant, C538 in the G220C and G221U mutants undergoes a ∼180° flip backward. In the A127G mutant, C538 only shows slight conformational movement, and in the hydrolysis reaction, G539 does not interact with G220. Instead, A127 assists C220 in the G220C mutant, and G127 helps G539 of A127G mutant, ultimately stabilizing the Pre structure. The G221U mutant does not provide any assisting forces for G539 or G220. G539 and C134 form a non-canonical Watson-Crick base pair with three hydrogen bonds, compared to the weaker interaction with only two hydrogen bonds in other mutants.

**Figure 4:**
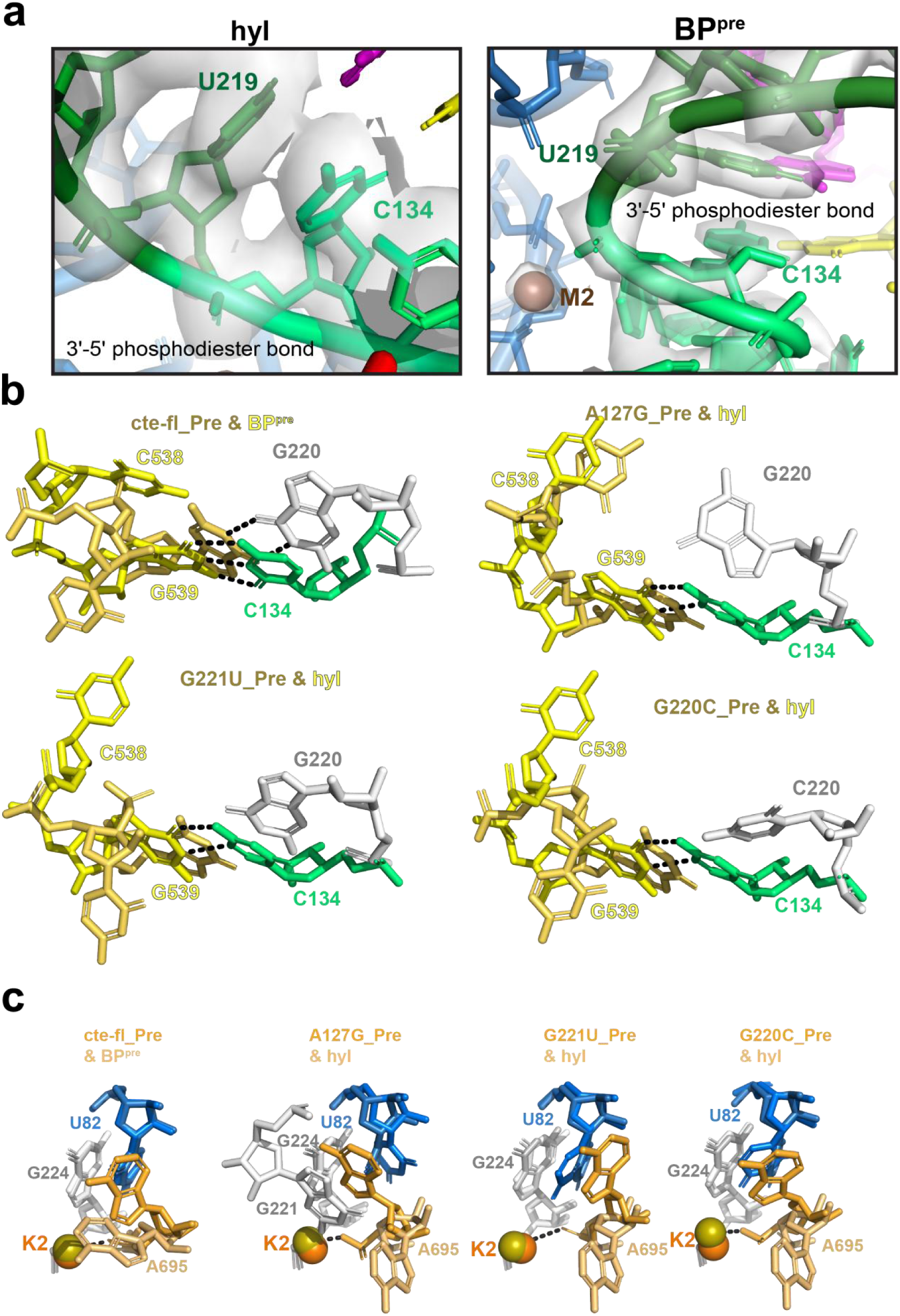
The conformational changes from 1^st^ step to 2^nd^ step by hydrolysis pathway. **a.** The zoom of circular RNA before forming 3*’*-5*’* phosphodiester bond in BP^pre^ state and formed in hyI conformation. **b.** C538 and G539 regulate the process from the 1^st^ step to the 2^nd^ step of the hydrolysis pathway in CP group II intron. **c.** A695 of J2/3 flips down from the 1^st^ step to the 2^nd^ step of the hydrolysis pathway.

During the transition from the 1^st^ to the 2^nd^ step of the hydrolysis pathway in the *O.i* Group IIC intron, G288 of J2/3 undergoes a ∼180° flip^12^. For *Cte* 1, the G696 of J2/3 base pairs with G61, and A697 base pairs with A60, forming specific G-G and A-A interactions (Fig. 4c). However, A695 undergoes a similarly significant ∼180° flip, accompanied by a conformational shift in U82 of the 2-nt bulge and a movement of G224, which have not been previously reported.

### The conformational changes from 2^nd^ step to hydrolysis product

Since we only obtained the hyI conformation from our cryo-EM samples of P1-E and D1–4 under CaCl_2_ conditions, we wondered if reintroducing P1–E and D1–4 into MgCl_2_ conditions would activate the second step of the reaction and allow us to capture the conformation of the hydrolysis products (movie S2). Analysis of the resulting circular RNAs revealed that the products from both the branching pathway and the direct second-step reaction of the hydrolysis pathway are identical, confirming that the correct reaction proceeded under MgCl_2_ conditions (Fig. 3b, Extended Data Fig. 11d, 11e). We subsequently analyzed cryo-EM sample of incubating P1–E with D1–4 under MgCl_2_ conditions, resulting in the identification of two hydrolysis product conformations, hyP1 and hyP2, in which the circularized exon has been released (Fig. 3b, 3g, 3h, Extended Data Fig. 10). By comparing the conformations of hyI and hyP1, we observed a significant conformational shift in the AUX-EBS (Extended Data Fig. 12a), and A695 flips back to a position similar to that in the Pre conformation. During this process, the conformational changes of U82 and G224 are minimal (Extended Data Fig. 12b).

### Alternative hydrolysis product conformation suggests undocking and docking of P1–D6

While the hyP1 conformation, with D6 pointing downward, is expected, we observed a new D6 conformation in hyP2 that resembles the Pre state, positioned across D1 (Fig. 3h, movie S3). Comparison between hyP1 and hyP2 structures reveals that the AUX-EBS region remains largely unchanged, with the exception of a substantial upward and downward flipping of D6 and minor conformational shifts in J2/3 (Extended Data Fig. 12c). HyP2 may not represent the typical conformation post the second step of hydrolysis, as achieving this state from the hyI conformation would require an energetically unfavorable reversal of the D6 conformation. Instead, we propose that hyP2 may result from P1–D6 undocking from D1–4 due to structural instability, and a new P1–D6 docking back into D1–4. In this scenario, P1–D6 product and the P1–E substrate competes for docking onto D1–4 (Fig. 3a, h, 5b).

### The density of the water molecules instead of 2*’*-OH of branch point

We hypothesize that the water molecule responsible for nucleophilic attack is located precisely at the original 2*’*-OH position of branch point A127. Due to limitations such as the absence of the D6 domain or insufficient map quality, such water molecules have not been identified in earlier studies, though their positions have been modeled^12,25^. In our G221U mutant structure, we observed for the first time that, as the 2*’*-OH of A127 moves away from the active center, its base engages with a water molecule, anchoring it at the original 2*’*-OH position of branching. This water molecule is thus poised to attack the 5*’*-splice site (Fig. 5a).

**Figure 5:**
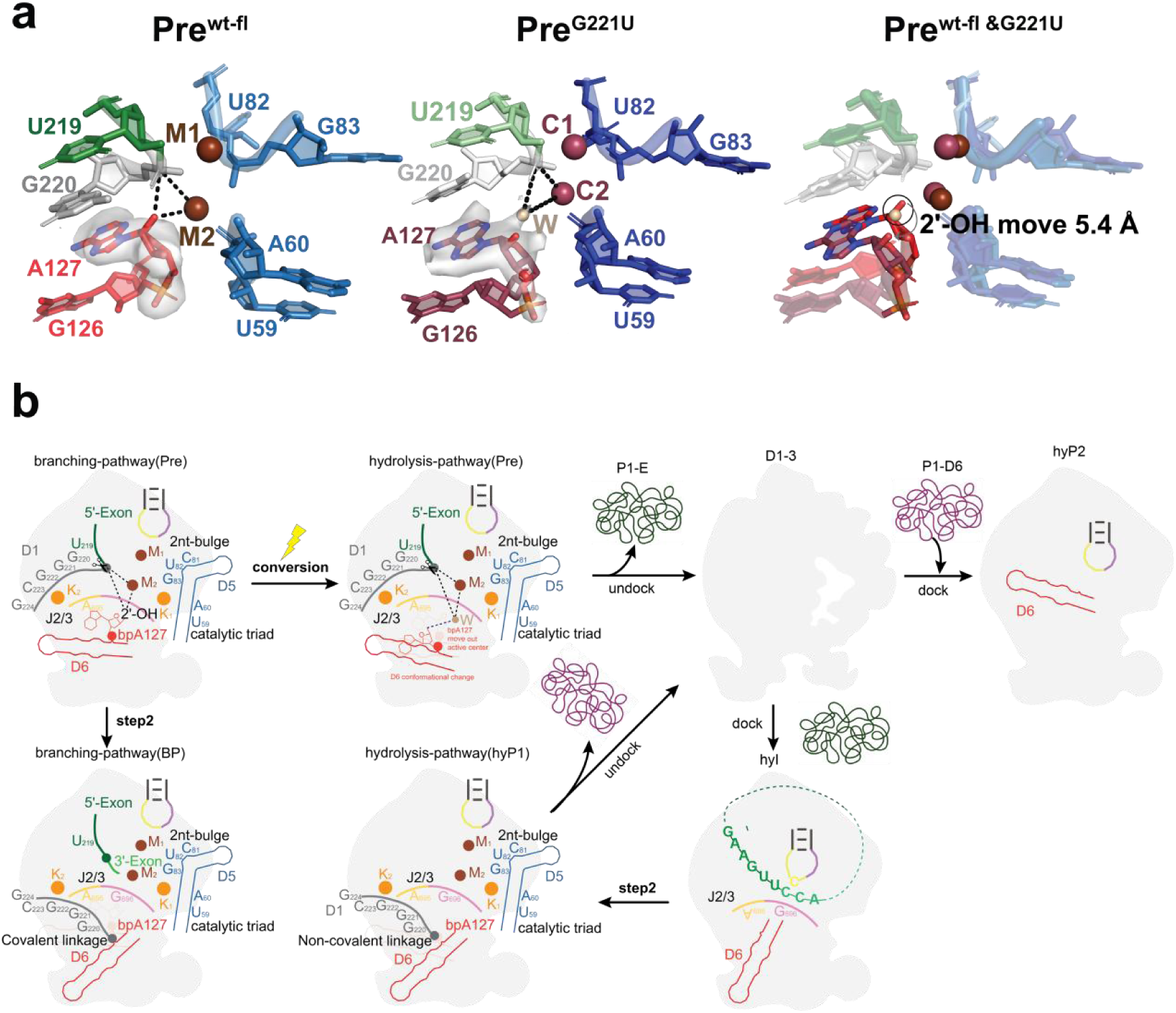
The mechanism of hydrolytic splicing. **a.** The water density of G221U compared with 2*’*-OH of A127 of wild type in Pre states. **b.** The mechanism of hydrolysis pathway.

## Discussion

### Life cycle of CP group II intron

Based on the structural information we have gathered, we propose a plausible life cycle of CP group II introns *in vivo*. First, upon transcription, the CP group II intron folds into a three-dimensional structure (the secondary structure diagram in the Extended Data Fig.13 was generated using the RNArtist software based on the already resolved tertiary structure) capable of performing back splicing. In the wild type, it transitions from the Pre conformation to the BI state through a change in the D6 conformation, completing two-step transesterification reactions that releases the circular RNA. This process occurs via the back-splicing branch pathway (Fig 5b, indicated by the blue arrows in the Extended Data Fig. 13).

In the presence of natural branch point A mutations or under stress conditions that generate branch-point or splice-site mutations, etc., the intron can follow the hydrolysis pathway. In the hydrolysis pathway, once the 1^st^ step is complete, the P1-Exon may proceed to the second step of splicing or undock from the D1–4 scaffold and dock back to initiate the second step, generating circular RNA (Fig 5b, indicated by the orange arrows in the Extended data Fig. 13). Although the undocking and docking of the P1-Exon after the first step of splicing is an energetically unfavorable process, it is a necessary event due to the requirement for circular RNA production by the CP group II intron or the spliceosome. The P1-Exon is inherently inverted, making this process inevitable. This undocking and docking of the P1-Exon during the generation of circular RNA via the hydrolysis pathway is a novel observation that adds to our understanding of the previously defined mechanism of linear exon splicing through the hydrolysis pathway, which had not been mentioned before. Since hydrolysis intermediates and hydrolysis products conformations without covalent bond formation, the P1-exon and P1-D6 of these products may be in a dynamic process of docking and undocking. As a result, there may also be a conformation of hydrolysis product2 in which D6 lies across D1. This conformation has never been observed before (Fig 5b, indicated by the orange arrows in the Extended data Fig. 13).

### Changes in D6, AUX and 5*’* SS in the branching-to-hydrolysis pathway transition

C538 of AUX in A127G mutant flips down to insert into a small groove formed by C100-G101-C102 and G126-G127 in D6, creating a triple interaction with G126 and G101 in D6. This results in the mutated G127 base pairing with C100, ultimately acting as a wedge that stabilizes the conformational changes (Extended Data Fig. 7). Moreover, G220 of the 5′ SS interacts with G539 in AUX, forming the important δ-δ′ interaction in the wild type, while it is involved in D6, base pairing with C102, which leads to a weak interaction between G221 and U103 (Extended Data Fig. 7).

In the G221U structure, C538 of AUX does not insert into the small groove of D6 (Extended Data Fig. 7), suggesting that the AUX conformation change is not necessary for the conversion to the complete hydrolysis pathway. The δ-δ’ interaction between G220 and G539 is also weakened, leading to a noticeable shift of 3.8 Å of D6 (Extended Data Figs. 7). The overall catalytic center of G221U mutant is similar to the wild type, with the most significant change being the displacement of D6. As the G221U mutant is also a complete hydrolysis mutant, a displacement of more than 3.8 Å in the D6 conformation appears sufficient for the transition from the branching pathway to the hydrolysis pathway.

The 5′ SS mutant G220C splices via both branching and hydrolysis pathways, with comparable intermediate products from each pathway (Fig. 1b and Extended Data Fig. 1c). G539 interacts with A127, while G221 forms two separate triple interactions with C102–U124 and G101–C125 (Fig. 2, Extended Data Fig. 7). Although the conformation of the 5′ SS undergoes significant changes, D6 conformation does not show substantial displacement (Extended Data Fig. 7b). In summary, significant shift in the D6 conformation is observed in the complete transition from the branching pathway to the hydrolysis pathway, whereas changes in AUX, as seen in the A127G, and in the 5′ SS, as observed in the G221U and G220C mutants, can also be involved in this transition process.

### Implications of “trans-splicing” of CP group II intron

Our structural study underscores the scaffolding function of D1–4, which enabled the “trans-splicing” between P1–E and D1–4. Our observation of an alternative post-2^nd^ step structure (hyP2) in this trans-splicing reaction suggests that P1–D6 may undock and dock back. We hypothesize that P1–E from other group II introns may also dock onto the non-self D1–4 scaffold to facilitate the generation of heterologous circular RNA (indicated by the gray arrows in the Extended Data Fig. 13).

At a first glance, the reconstructed 2^nd^ step of the hydrolytic splicing involving P1–Exon and D1–4 to produce RNA circles resembles trans-splicing^26,27^, where exons from two separate molecules are spliced together. This type of splicing occurs naturally in spliceosome^28^, group I intron^29^, or group II intron^30^ based splicing, and can be engineered for therapeutics^31^. However, our strategy is distinct for several reasons. First, a typical trans-splicing undergoes two steps as in conventional splicing, whereas our strategy only involves the second step of splicing, bypassing the first step. Second, in a typical trans-splicing the ligated exons come from two separate molecules, whereas in our strategy the exons reside in one molecule in an inverted manner to produce circles. Nevertheless, both typical trans-splicing and our finding highlight the modular nature of group II intron, for which we now provide a structural basis. In CP group II intron, D1–4 acts primarily as a stable “scaffold module”, to which the “catalytic module” P1–D6 can dock back, bringing in the inverted exon and catalyzing circRNA formation. From an evolutionary standpoint, trans-splicing is thought to facilitate exon shuffling and gene evolution. Based on our study of the CP group II intron, we speculate that intron modules from different sources/origins may be able to form a functional ribozyme in trans^21^, and this mix-and-match might facilitate the evolution of group II intron themselves, eventually leading to the modern spliceosome. It would be interesting to search for such orphan group II intron modules in diverse genomes. Indeed, disrupted group II introns have been reported^32–37^, with extreme examples in which the intron is split into at least three pieces^33,38–40^. From an engineering perspective, our finding provides a streamlined strategy to produce circular RNA, with a constant scaffold module and variable catalytic-exon module.

### Implications of the hydrolysis pathway

Structures of hydrolytic splicing have now been shown for group IIC introns^12^, reverse splicing of HYER^25^ (Extended Data Fig. 11a–c), and back splicing of CP group II, indicating that hydrolysis pathways occur in various splicing processes. It has also been suggested that the hydrolysis pathway may be ancestral to the branch pathway^8^. Given that group II introns could also be ancestral to spliceosomes^41^, different splicing forms have been reported to have an evolutionary relationship with spliceosome structure and mechanisms^42^. Although it is currently believed that the major spliceosome in humans does not utilize a hydrolysis pathway^43^, the second step of the hydrolysis pathway in CP group II introns resembles the second step of trans-splicing, which has been extensively documented in eukaryotes and humans. Therefore, it is possible that other spliceosomes in humans are capable of following hydrolysis pathways to generate circular RNA or linear RNA. Our current findings underscore the complexity and evolutionary significance of splicing mechanisms, highlighting the need for further investigation into the functional diversity of intron splicing pathways.

## Materials and Methods

### RNA preparation

The *Cte* group II intron sequence was derived from NCBI (GenBank: NZ_AAUJ02000001.1). The wild type and mutated RNAs were prepared as described previously^44^. In brief, the DNA templates in pUC-57 plasmid were amplified with the regular forward primer and reverse primer (supplementary excel). In vitro transcription of all RNAs in this article were carried out with 0.3 µM DNA template, 5 mM NTPs and 1 U/µL RNase inhibitor in 1× transcription buffer containing 50 mM Tris-HCl (pH 7.9), 0.01% TritonX-100, 20 mM MgCl_2_, 2 mM spermidine, 10 mM DTT, and incubated at 37 °C for 1 h^45^. The transcription products were mixed with 2× denaturing gel loading buffer containing 95% formamide, 0.025% SDS, 10 mM EDTA, 0.025% xylene cyanol, and 0.025% bromophenol blue, and loaded on an 4% 19:1 acrylamide:bis, 8 M urea polyacrylamide gel. The gel was run at 15 W for 3 h, then visualized briefly with a 254-nm UV lamp, held far from the gel to minimize RNA damage^46^. Then the RNA was eluted from the gel overnight in elution buffer containing 30 mM sodium acetate (pH 5.2) and 5 mM EDTA on an active rotator at 4 °C. The resulting gel slurry was then filtered through 0.45 μm filters (Minisart® Syringe Filter, Sartorius). The resulting RNA was and precipitated with equal volume of isopropanol to remove urea and salts, then washed by 75% ethanol for 2-3 times. Air-dry the pellet and resuspend with RNase-free water. The RNA was quantified using NanoDrop spectrophotometer (Thermo Scientific) and kept at -80 °C before use.

### Splicing reactions

Splicing reactions of 0.05 μM wild-type and mutated *Cte*RNAs were carried out in splicing buffer 40 mM HEPES (pH 7.5), 240 mM KCl at 23 °C. The RNAs were heated to 80 °C for 1 min and cooled to 60 °C (0.5 °C/s) in the splicing buffer. After annealing, a final concentration of 20 mM or 100 mM MgCl_2_ was added to start the splicing assay, samples were then incubated at 23 °C for 30 min.

For cryo-EM sample preparation, the splicing reaction was activated in a total volume of 0.5-1 ml as described above. The mixture was then rapidly chilled on ice and concentrated to approximately 15 μM by concentrator columns with 100 kDa cutoff (Ultrafiltration Centrifugal Tube, Millipore) at 4 °C.

For subsequent biochemical mechanism investigations, the splicing assay was quenched by adding an equal volume of denaturing loading buffer containing 100 mM EDTA, 8 M urea, 0.1% SDS, 0.025% xylene cyanol and 0.025% bromophenol blue. Then the electrophoreses were performed on denaturing 8M urea and 6% 19:1 acrylamide:bis gel, running at 12 W for 2 h. The gels were stained for 10 min in SyBrGold (Invitrogen) and visualized by Typhoon FLA 9500 (GE).

### Cryo-EM sample preparation

A total of 3 μl of the resulting splicing products (about 15 μM) was applied onto glow-discharged (10 s) 200-mesh R1.2/1.3-2nmC Quantifoil Cu grids. The grids were blotted for 1.5 s in 100% humidity at 4 °C with no blotting offset and rapidly frozen in liquid ethane using a Vitrobot Mark IV (Thermo Fisher).

### Cryo-EM data collection and image processing

Cryo-EM data were acquired using a Titan Krios microscope and Krios G4 (Thermo Fisher) operating at 300 kV and 200 kV. The Titan Krios microscope was equipped with a K3 summit direct detector (Gatan) and a GIF quantum energy filter (Gatan) set to a slit width of 20 eV. The Titan Krios microscope was equipped with a Falcon 4i summit direct detector (Thermo Fisher). Automated data collection was performed using the EPU software^47^ and SerialEM software^48^ with the beam-image shift method^49^.

Movies of the Cte-fl_MgCl_2_, A127G_MgCl_2_ were captured in super-resolution mode at a nominal magnification of 100,000× for Cte-fl_MgCl_2_ and 105,000× for A127G_MgCl_2_ resulting in a physical pixel size of 0.959Å for Cte-fl_MgCl_2_ and 0.832 Å for A127G_MgCl_2_. Movies of the U82G_CaCl_2_, G220C_CaCl_2_, G221U_CaCl_2_, trans-splicing_CaCl_2_ and trans-splicing_MgCl_2_ were separately recorded at a magnification of 120,000×, corresponding to a physical pixel size of 1.2 Å. Movies of the U133G_CaCl_2_, were recorded at a magnification of 100,000×, corresponding to a physical pixel size of 0.82 Å. Each movie covered a defocus range from -1.2 μm to -2.2 μm. Movie stacks were dose-fractionated into 40 frames, with a total exposure dose of approximately 50-51 e-/Å^2^.

In cryo-EM data processing, a standardized procedure was employed throughout the analysis within cryoSPARC v4^50^. The initial steps involved patch motion correction and patch contrast transfer function (CTF) estimation to rectify stage shift, beam-induced motion, and imaging distortions. Rigorous criteria were applied during the selection of high-quality images, taking into account factors such as ice thickness, defocus range, and estimated resolution.

The particle processing pipeline began with initial particle picking using blob picking and template picking, followed by two rounds of reference-free 2D classification. Subsequently, a subset of high-quality particles underwent ab-initio reconstruction, yielding initial references. All viable particles then underwent multi-round 3D classification without mask. Classes containing well-defined D6 domains were re-extracted without binning. Through non-uniform refinement, local motion correction, and global refinement without mask and local refinement with mask, high-resolution maps were obtained. The reported resolutions above are based on the gold-standard Fourier shell correlation (FSC) 0.143 criterion. Subsequently, local resolution estimation was conducted. The final refined maps underwent separate sharpening using either DeepEMhancer^51^ or EMReady^52^. The resulting sharpened maps, featuring enhanced density, were employed for subsequent structural analyses. All the visualization and evaluation of 3D density maps were performed with UCSF Chimera^53^ and ChimeraX^54^. Detailed cryoEM data processing workflows are outlined as indicated in Fig. S2, S4, S5, S6, S8 and S9.

### Cryo-EM model building and refinement

We de novo reconstructed the Cte-fl_Pre model by the information of IBS-EBS according to the conserved secondary structure as the starting point of model. The other has no high resolution of map manually adjusted and rebuilt with Coot^55^ according the Alphafold3 prediction model. These models were refined with Phenix.real_space_refine^56^, yielding an averaged model-map correlation coefficient (CCmask) in range from 0.84 to 0.93, respectively. Final models were validated by MolProbity^57^. Secondary structure diagrams were drawn with RNArtist then manually regulated at Adobe Illustrator.

## Acknowledgements

Cryo-EM data were collected at the Center of Cryo-Electron Microscopy at Life Medicine in Fudan University and the Cryo-EM Core Facility at UMass Chan Medical School. The data were processed at Life Medicine High Performance Computing Center in Fudan University and HPC cluster in UMass Chan Medical School. We thank Dr. Hongyun Shen from the Center of Cryo-Electron Microscopy at Life Medicine in Fudan University for helping to collect 300 kV cryo-EM data and Kangkang Song from the Cryo-EM core facility at UMass Chan Medical School for helping to collect 200 kV and 300 kV cryo-EM data. This work was supported by Natural Science Foundation of China (NSFC 31971130) to J.M.

## Author contributions

X.L. conceived the project; J.M. and Z.C. supervised the project; X.L. and Y.Y. prepared cryo-EM samples and performed electrophoresis and analyzed gel electrophoresis results; X.L., K.S., C.X. and Y.Y. collected cryo-EM data; X.L. and Y.Y. processed cryo-EM data. X.L, G.D., J.G. and J.M. built, refined and validated atomic coordinate models. X.L. and Y.Y. prepared the initial manuscript. Other authors contributed to the preparation of the manuscript.

## Data availability

The cryo-EM maps and associated atomic coordinate models of Cte-fl_MgCl_2__Pre, Cte-fl_MgCl_2__BP^pre^, A127G_MgCl_2__Pre, A127G_MgCl_2__D1-3, G220C_CaCl_2__Pre, G221U_CaCl_2__Pre, hydrolysis-intermediate_CaCl_2_, hydrolysis-product1_MgCl_2_ and hydrolysis-product2_MgCl_2_ have been deposited in the wwPDB OneDrop System under EMD accession codes EMD-74859, EMD-74867, EMD-74857, EMD-74868, EMD-74869, EMD-74871, EMD-74872, EMD-74877, EMD-74874 and PDB ID codes 9ZV6, 9ZVB, 9ZV5, 9ZVC, 9ZVD, 9ZVF, 9ZVG, 9ZVK, 9ZVI respectively.

## Supplement Figures

**Extended Data Fig. 1:**
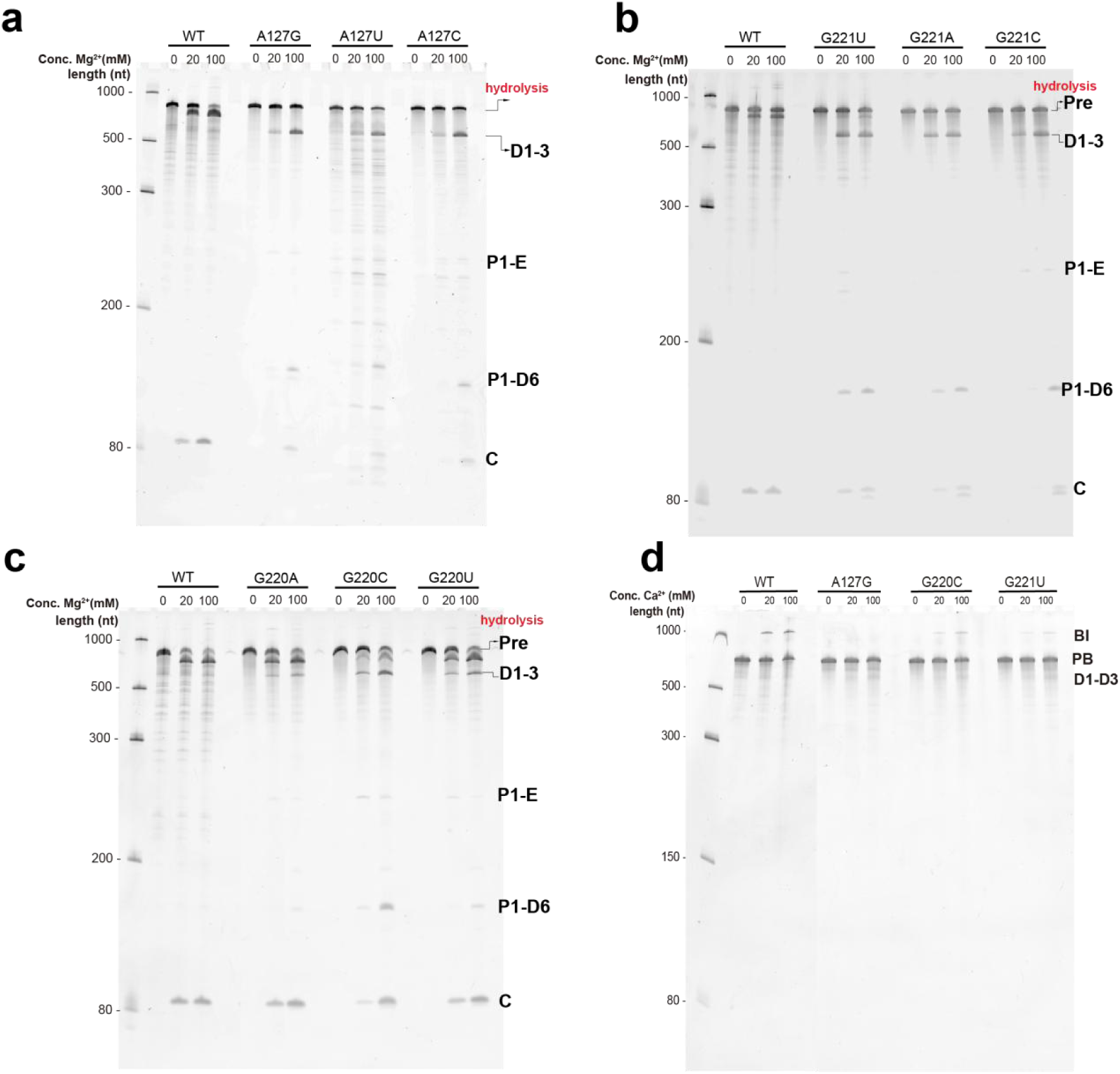
Biochemical analyses of *Cte* 1 wild type and mutants. The back-splicing activity of *Cte* 1 hydrolysis mutants was analyzed at different Mg²⁺ (**a-c**) and Ca^2+^ (**d**) concentration. The G220C and G221U samples at 20 mM CaCl_2_ condition were for cryo-EM. The experiments shown in these panels were repeated more than three times with similar results.

**Extended Data Fig. 2:**
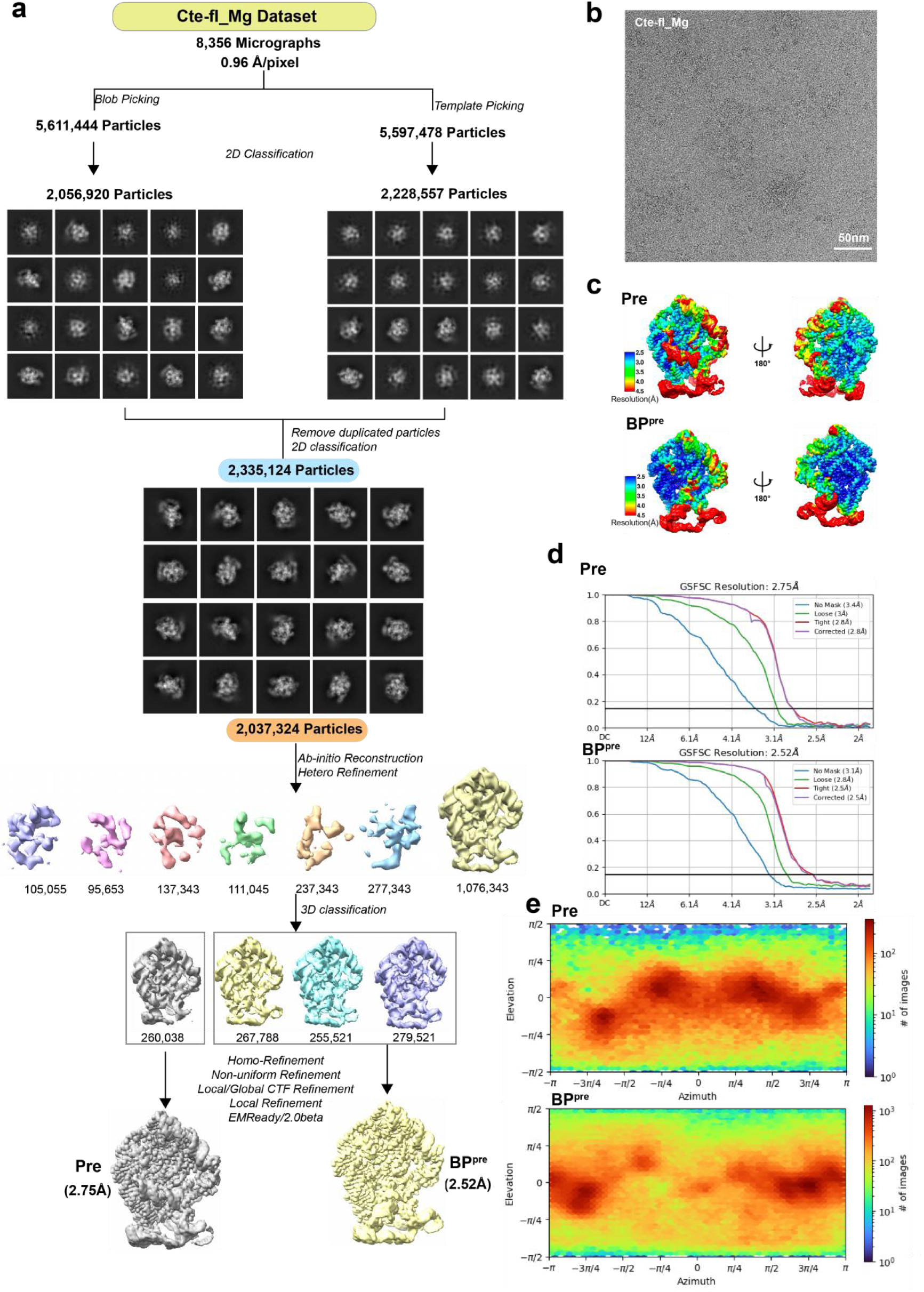
Cryo-EM workflow of the wild type *Cte* 1 full length in MgCl_2_ condition. **a.** Cryo-EM data processing workflow of the wild type *Cte* 1 full length at the MgCl_2_ condition. **b-e.** The particle distribution of full micrograph (**b**) and local resolution map (**c**) and FSC curves with the gold standard threshold of 0.143 (**d**) and euler angular distribution (**e**) for *Cte* 1 full length Pre and BI conformation maps.

**Extended Data Fig. 3:**
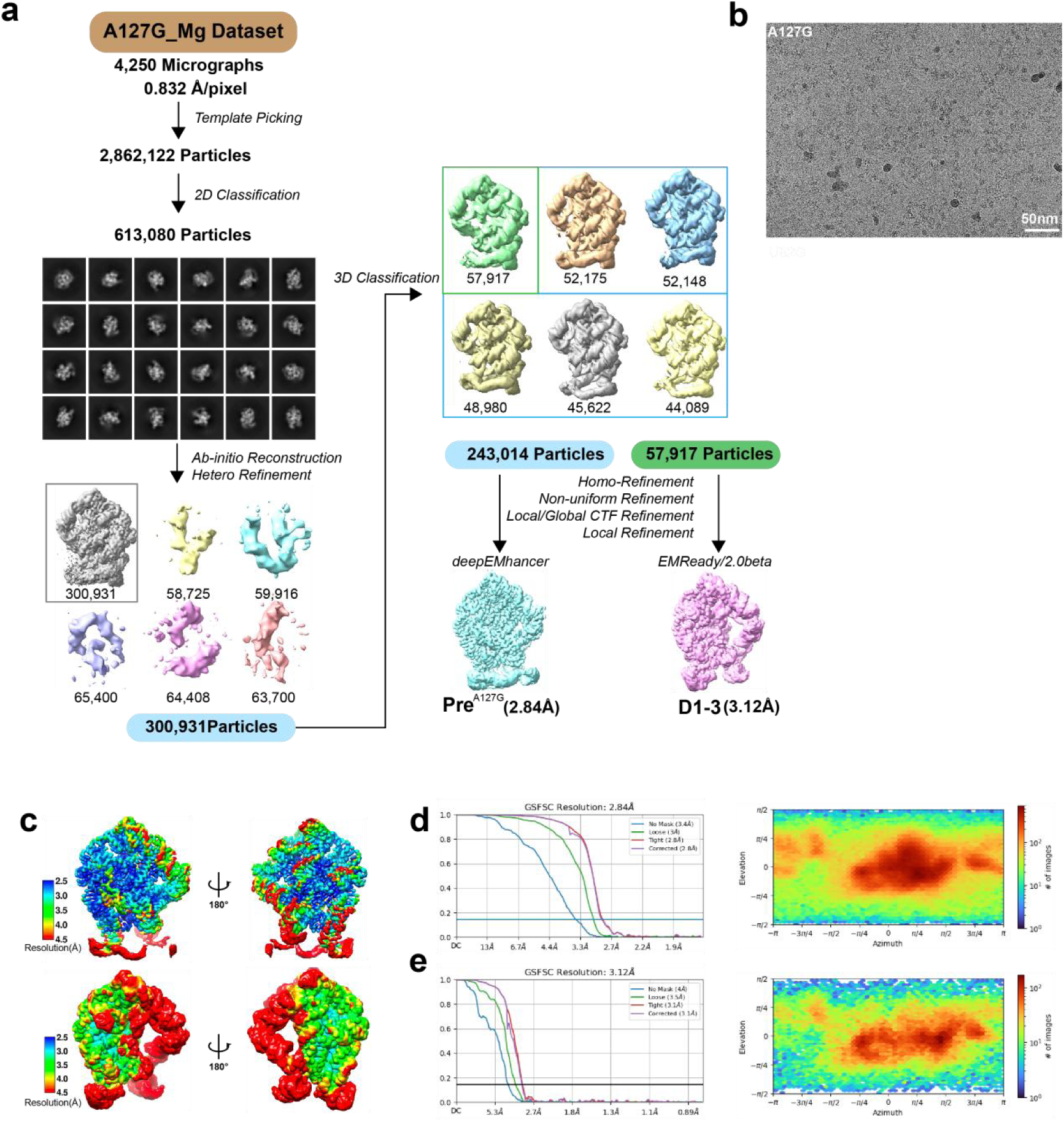
Cryo-EM workflow of the branch point A127G mutant in MgCl_2_ condition. **a.** Cryo-EM data processing workflow of the branch point A127G mutant sample at the MgCl_2_ condition. **b-e.** The particle distribution of full micrograph (b) and local resolution map (c) and FSC curves with the gold standard threshold of 0.143 (d) and euler angular distribution (e) for conformation maps.

**Extended Data Fig. 4:**
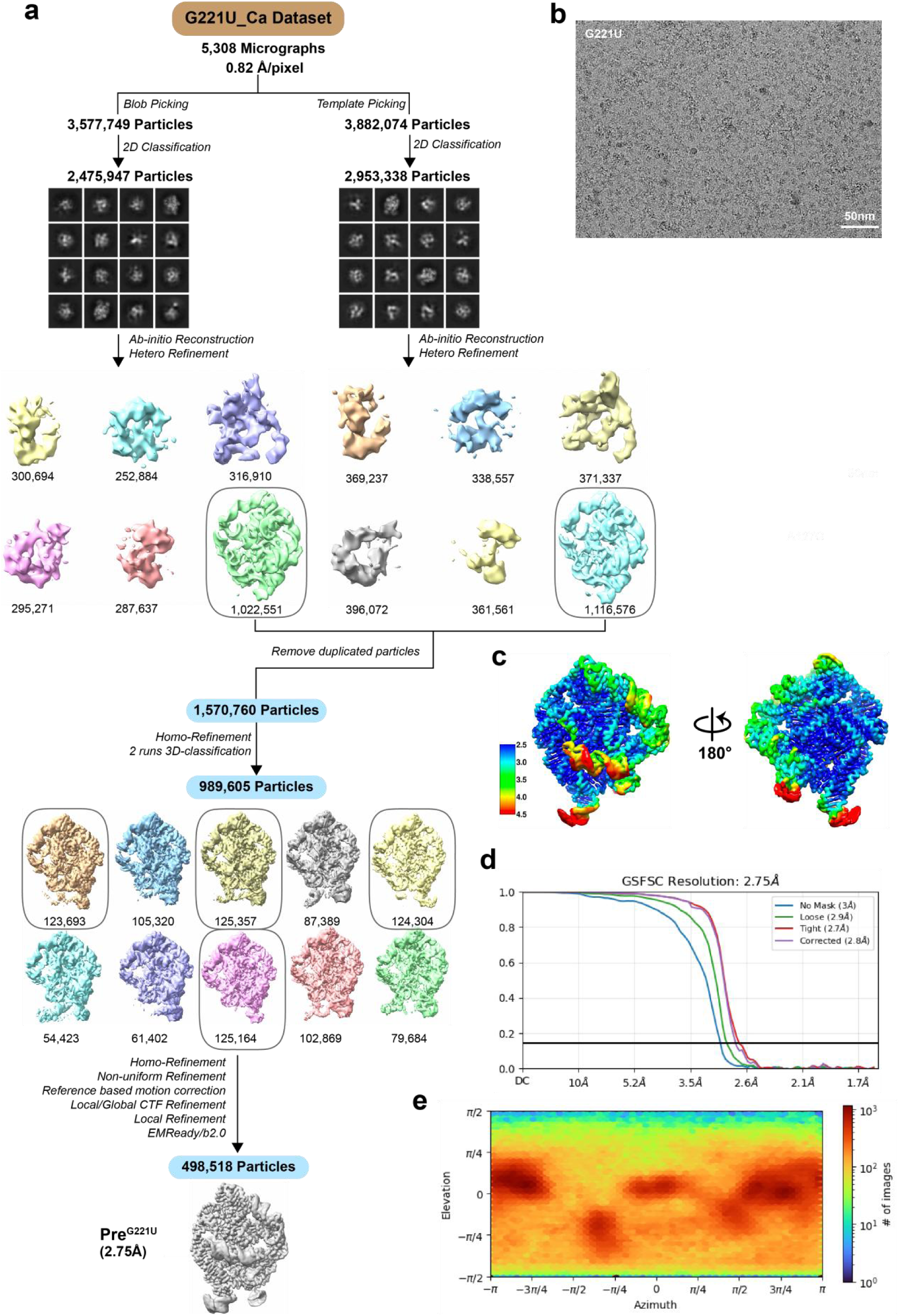
Cryo-EM workflow of the 5*’* SS mutant G221U in CaCl_2_ conditions. **a.** Cryo-EM data processing workflow of the 5*’* SS mutant G221U at the CaCl_2_ condition. **b-e.** The particle distribution of full micrograph (**b**) and local resolution map (**c**) and FSC curves with the gold standard threshold of 0.143 (**d**) and euler angular distribution (**e**) for conformation maps.

**Extended Data Fig. 5:**
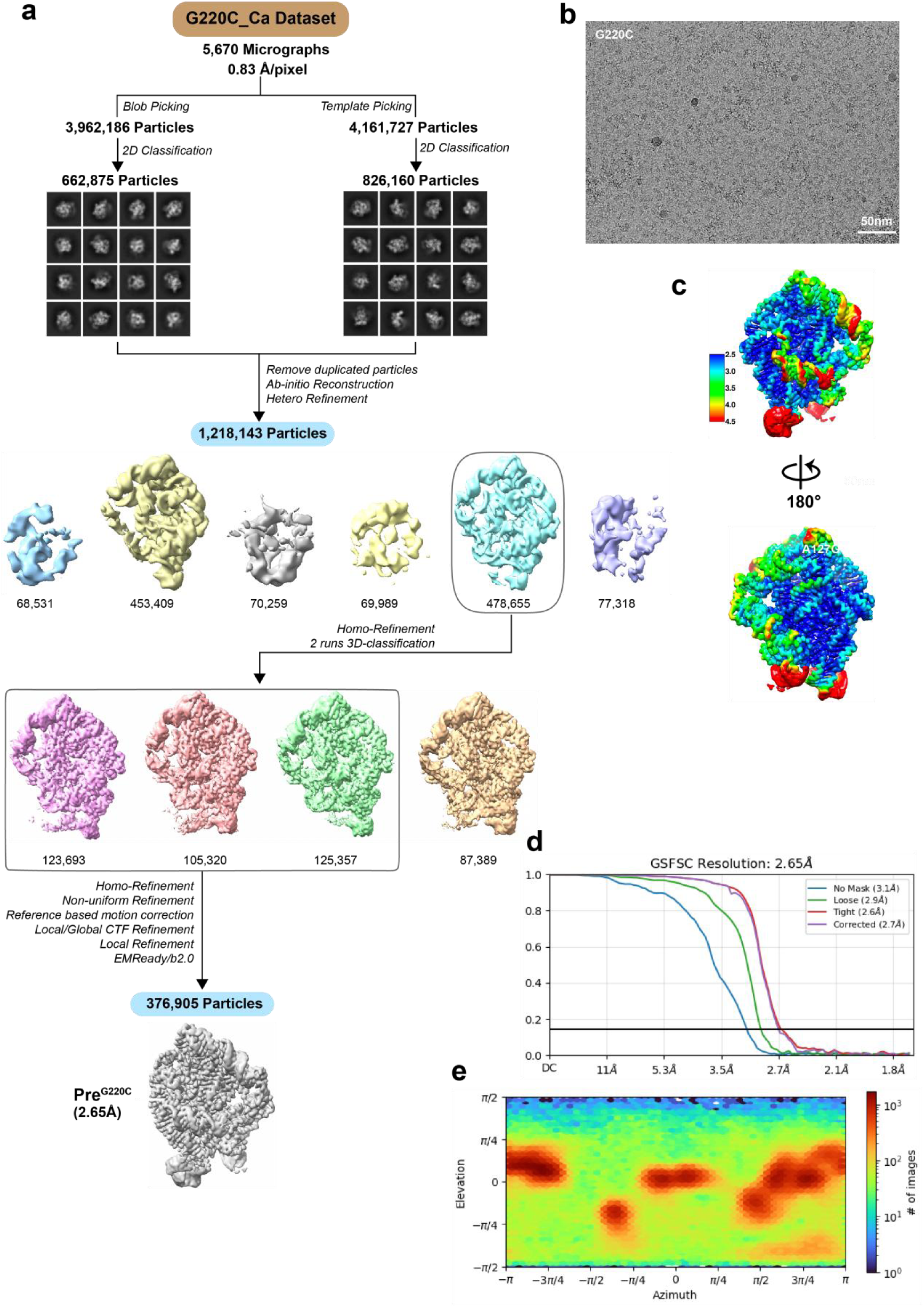
Cryo-EM workflow of the 5*’* SS mutant G220C in CaCl_2_ conditions. **a.** Cryo-EM data processing workflow of the 5*’* SS mutant G220C at the CaCl_2_ condition. **b-e.** The particle distribution of full micrograph (**b**) and local resolution map (**c**) and FSC curves with the gold standard threshold of 0.143 (**d**) and euler angular distribution (**e**) for conformation maps.

**Extended data Fig. 6:**
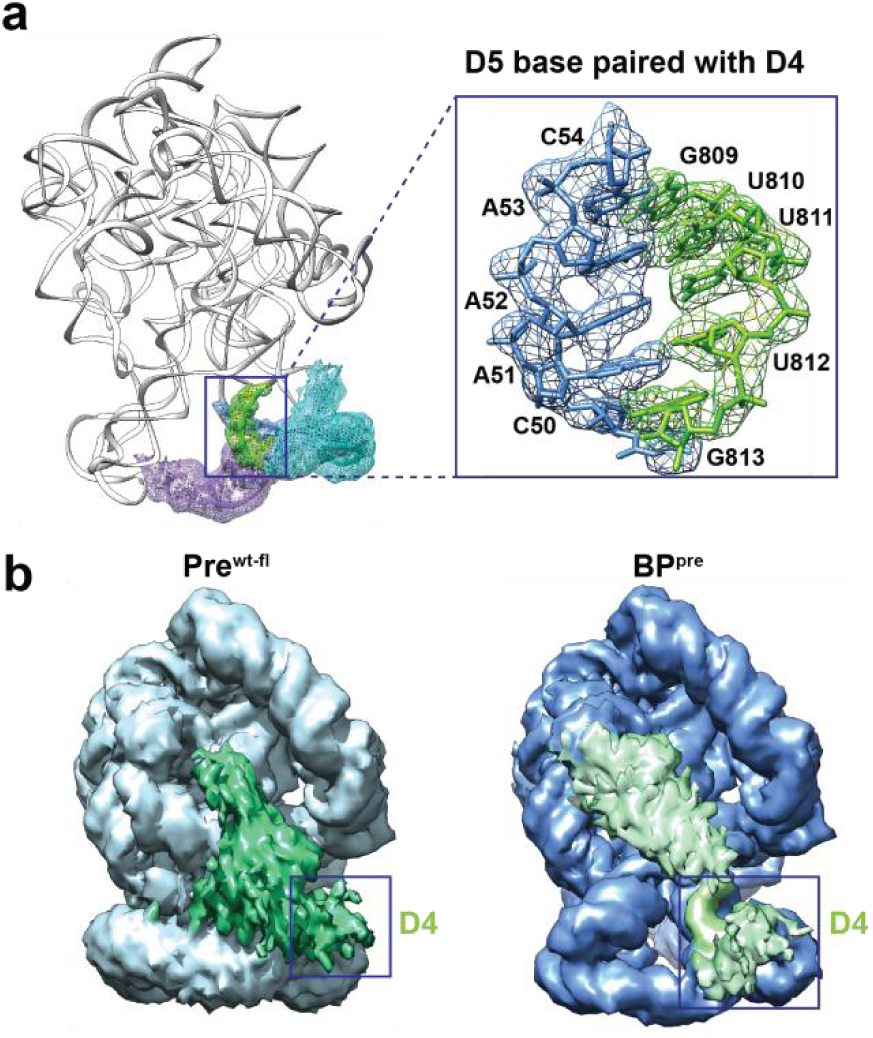
The extra density of D4 domain in *Cte* 1 wild type. **a.** The residues 809–813 of D4 are stabilized by base pairing with residues 50–54 of D5. **b.** The low density shows the D4 domain position of Pre^wt-fl^ and BP^pre^ states.

**Extended Data Fig. 7:**
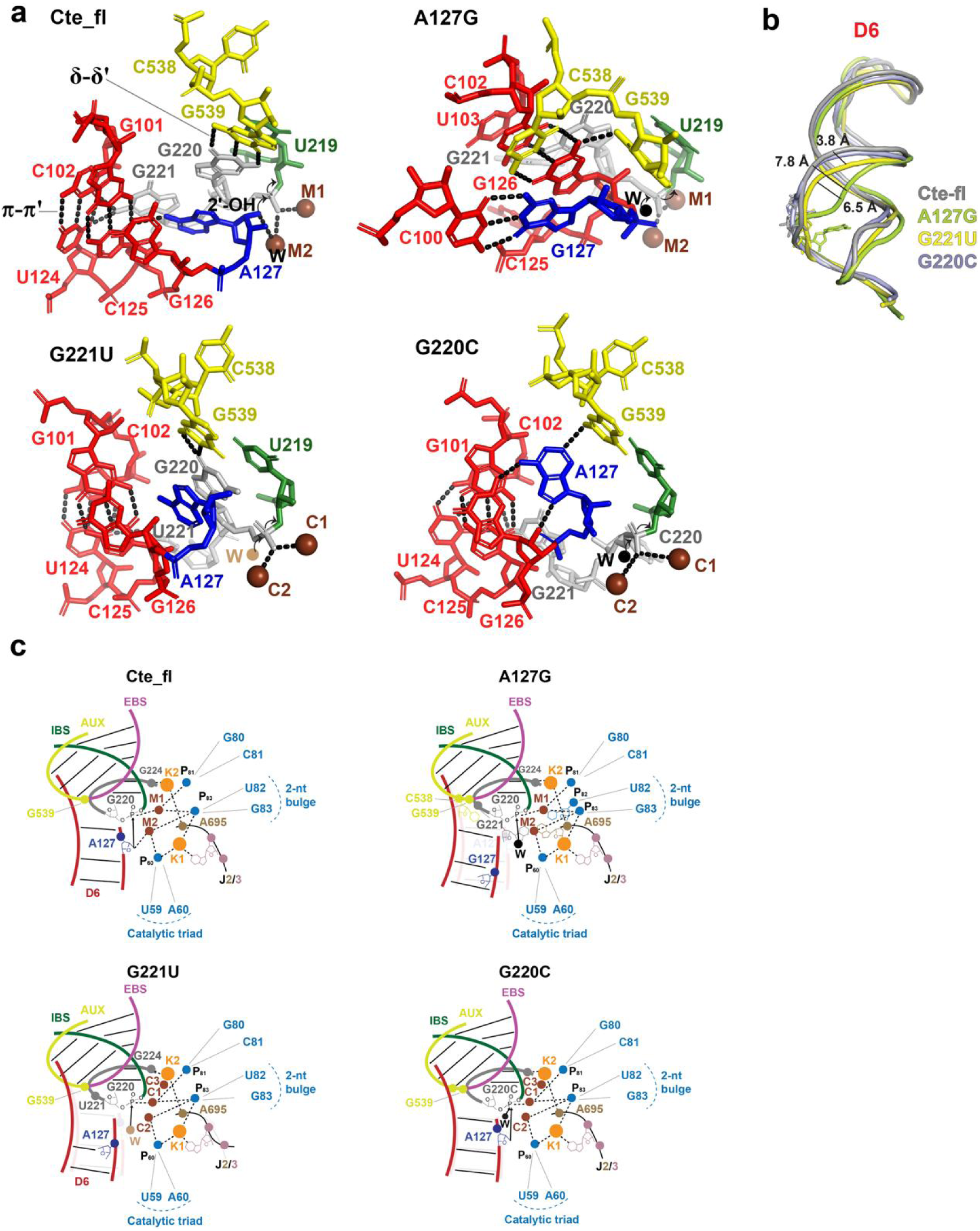
Comparison of the catalytic center between wild type and hydrolysis mutants. **a.** C538 of mutant A127G sticks into D6 small groove in the hydrolysis pathway. **b.** Schematic diagram of the catalytic center of *Cte* full length and hydrolysis mutants.

**Extended Data Fig. 8:**
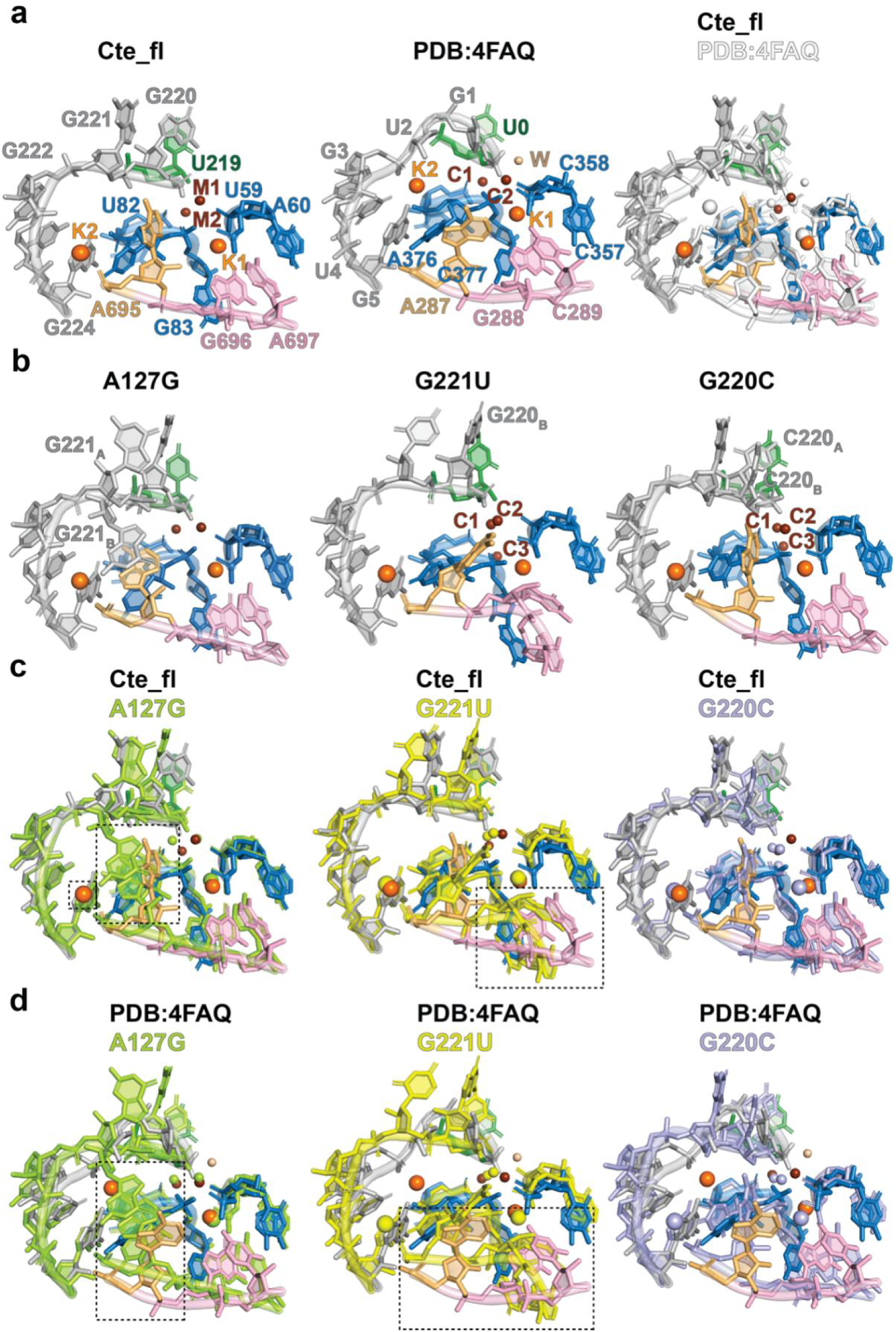
The triple helix conformation and mental cluster of hydrolysis pathway mutants in the Pre states. **a.** The triple helix conformation and mental cluster comparison of *Cte* full length (Cte-fl) and group IIC intron (PDB: 4FAQ) from *Oceanobacillus iheyensis*. **b.** The triple helix conformation and mental cluster of mutants A127G, G221U and G220C. **c-d.** The triple helix conformation and mental cluster comparison of Cte-fl or group IIC intron and mutants A127G, G221U and G220C.

**Extended Data Fig. 9:**
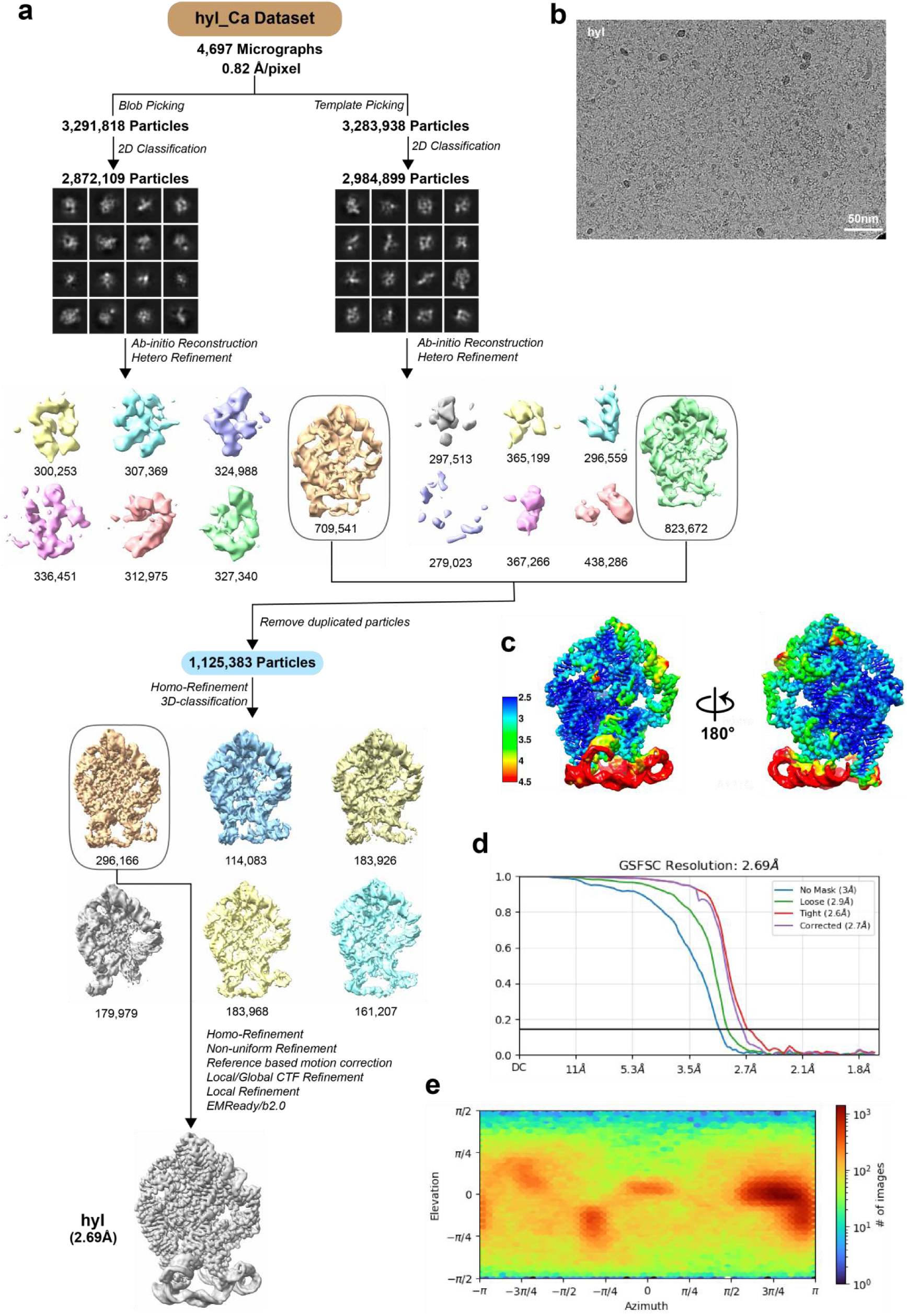
Cryo-EM workflow of the hydrolysis intermediate in CaCl_2_ conditions. **a.** Cryo-EM data processing workflow of the hydrolysis intermediate at the CaCl_2_ condition. **b-e.** The particle distribution of full micrograph (**b**) and local resolution map (**c**) and FSC curves with the gold standard threshold of 0.143 (**d**) and euler angular distribution (**e**) for conformation maps.

**Extended Data Fig. 10:**
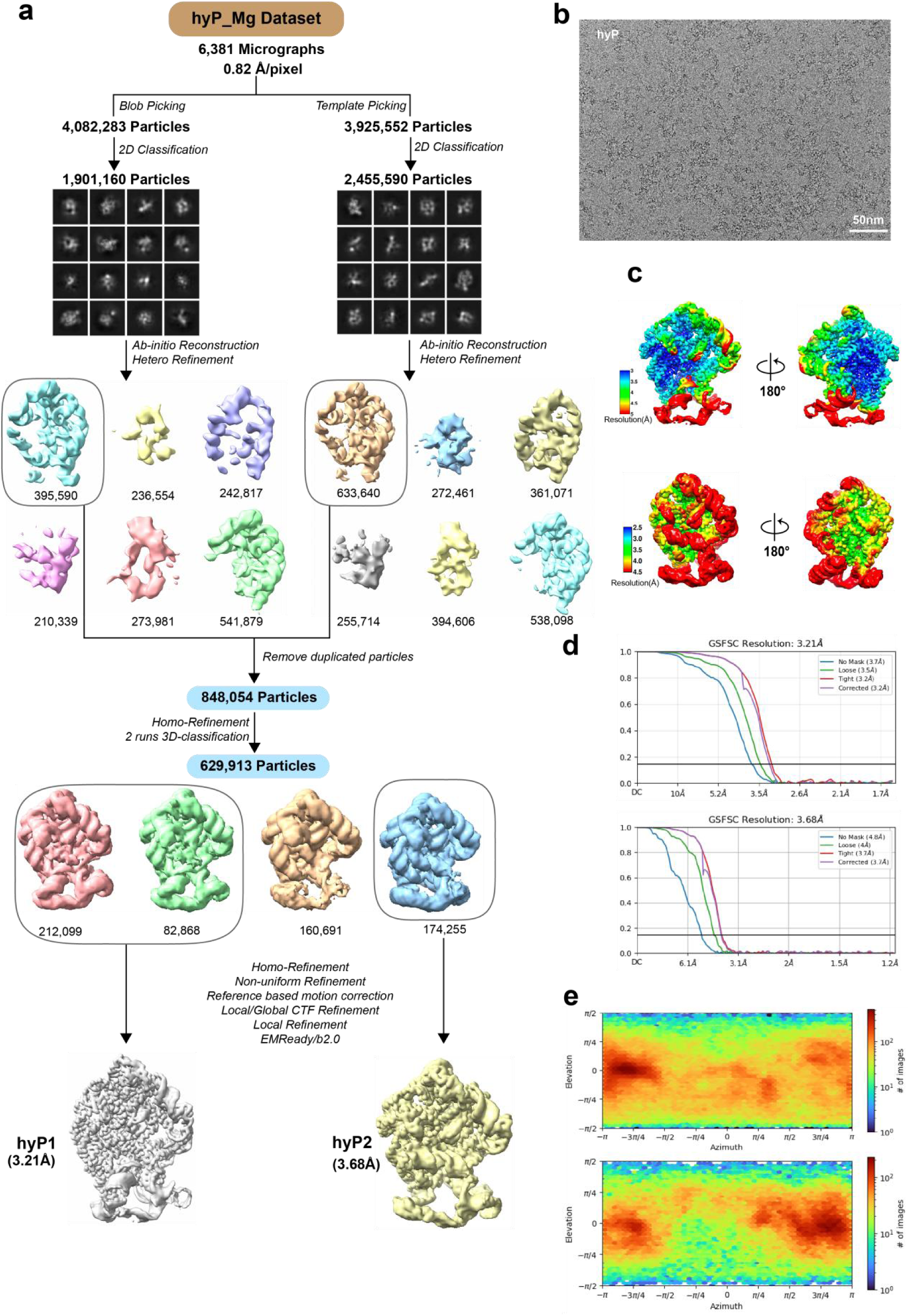
Cryo-EM workflow of the hydrolysis product in MgCl_2_ conditions. **a.** Cryo-EM data processing workflow of the hydrolysis product at the MgCl_2_ condition. **b-e.** The particle distribution of full micrograph (**b**) and local resolution map (**c**) and FSC curves with the gold standard threshold of 0.143 (**d**) and euler angular distribution (**e**) for conformation maps.

**Extended Data Fig. 11:**
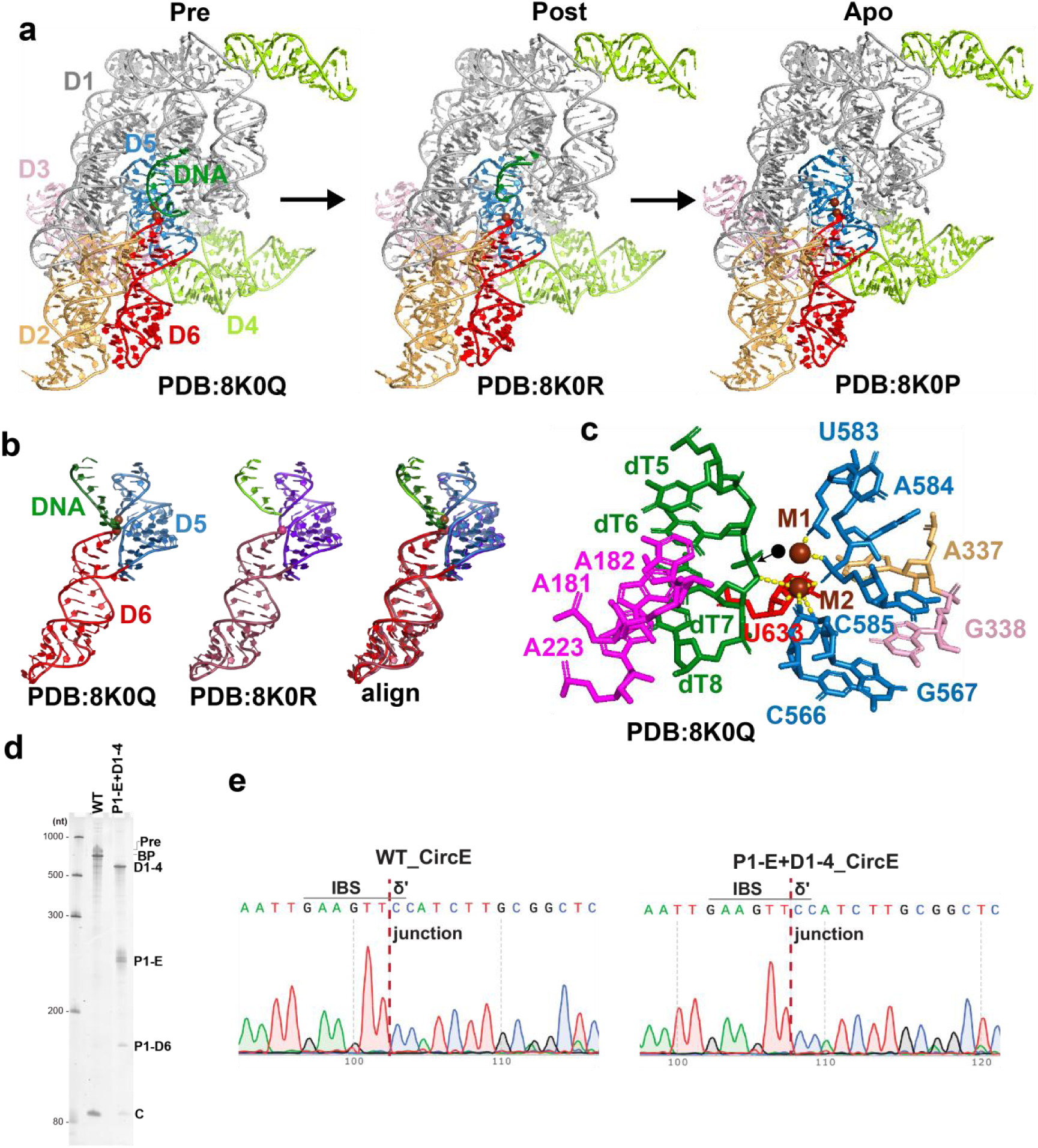
The catalytic center analysis of group IIC intron by hydrolysis pathway. **a.** The structures of group IIC intron HYER editing DNA substrate by the hydrolysis pathway in reverse splicing. **b.** The zoom in of D5-D6-Exon domain of HYER from pre state to post state and alignment. **c.** The catalytic center of HYER pre state. The black circle represents the theoretical position of water molecules. **d-e.** The Syber Gold-stained 6% urea–polyacrylamide gel electrophoresis (PAGE) and Sanger sequencing analysis of the CircE band on the gel show that *Cte 1* wild type and hydrolysis “trans-splicing” (P1-E+D1-4) produced expected circular exons (CircE) in 100 mM MgCl_2_ conditions.

**Extended Data Fig. 12:**
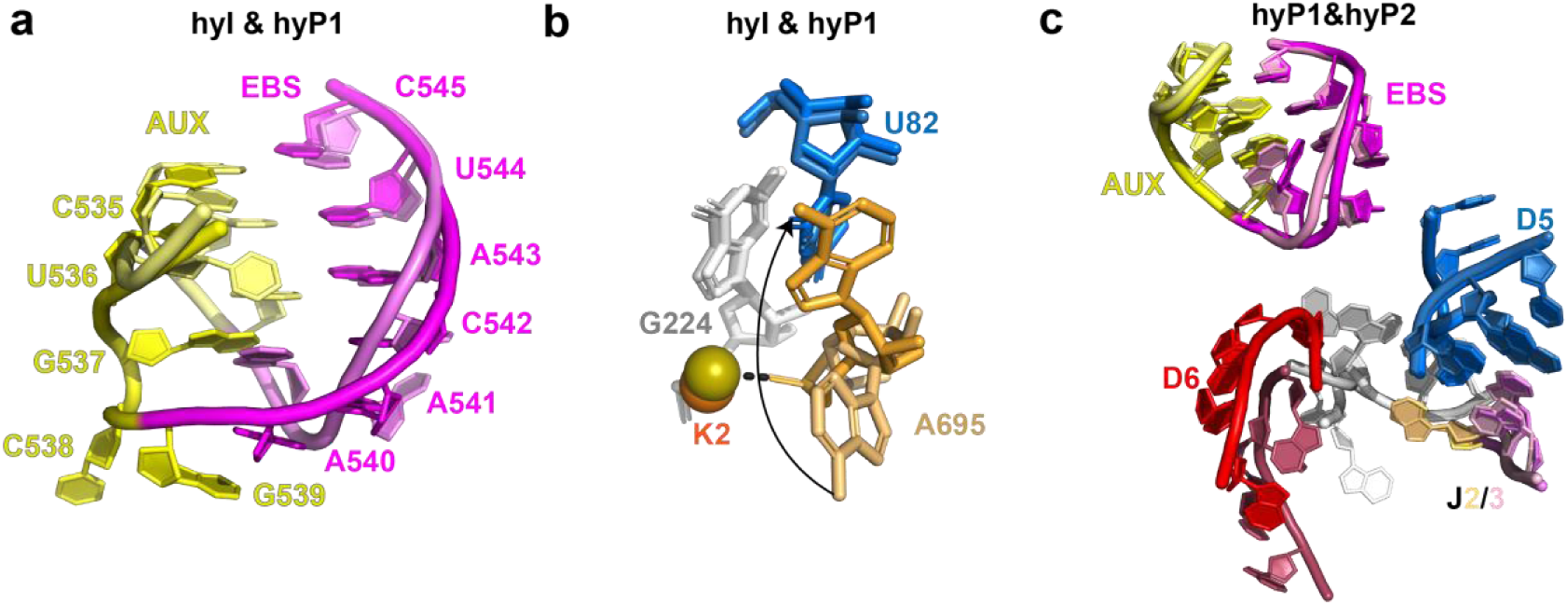
Analysis of the conformational changes from the 2^nd^ step to product release of CP group II intron in the hydrolysis pathway. **a.** AUX and EBS conformational changes from the hydrolysis intermediate (hyI) to the hydrolysis product1 (hyP1) states. **b.** A695 of J2/3 flips back from hyI to hyP1 state. **c.** The catalytic center alignment of hyP1 and hydrolysis product2 (hyP2) conformations.

**Extended Data Fig. 13:**
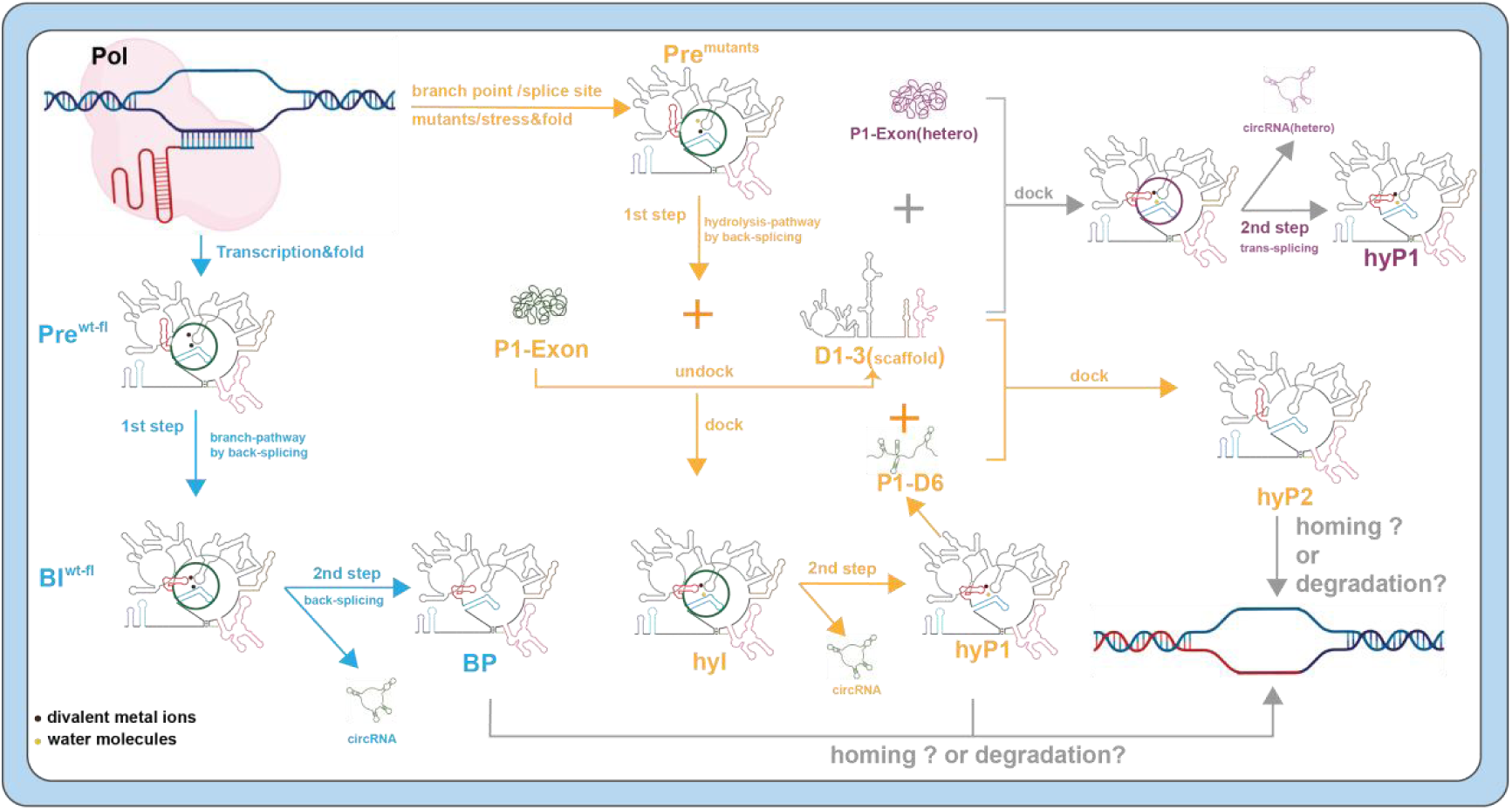
Life cycle of CP group II intron in vivo.

**Table S1:**
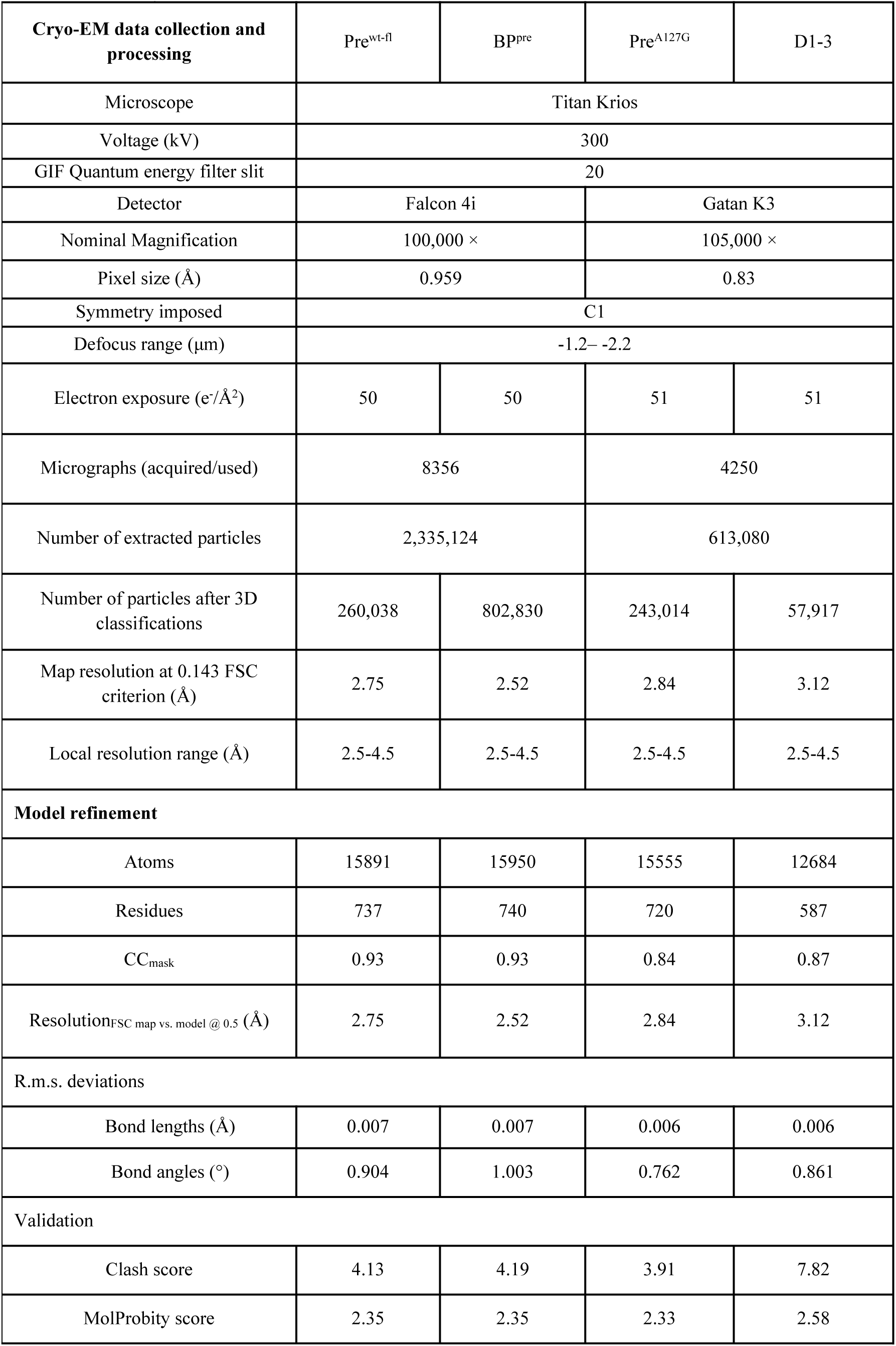
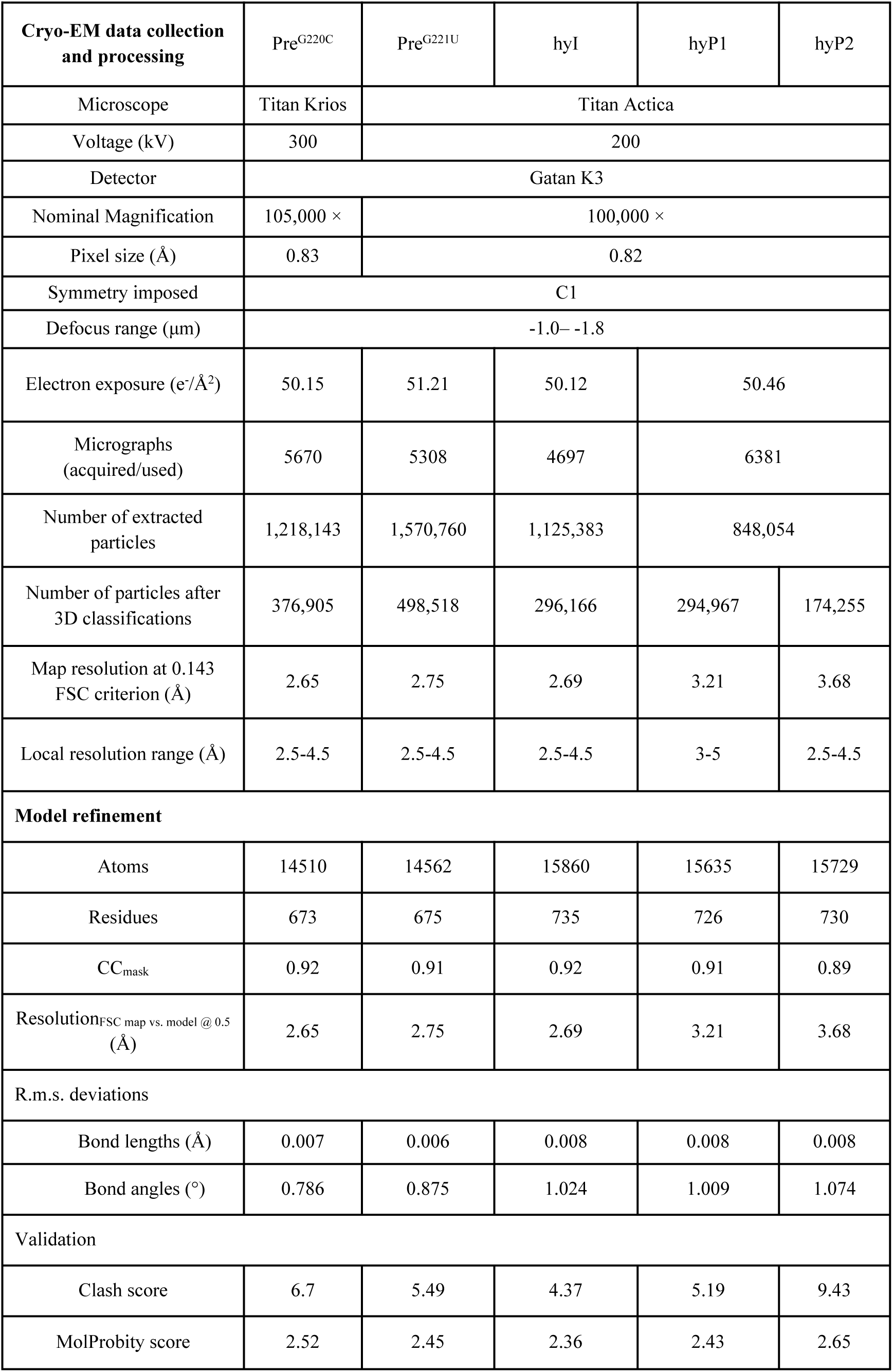
Cryo-EM Data Collection, Refinement and Validation Statistics

|  |  |  |  |  |
| --- | --- | --- | --- | --- |
| Cryo-EM data collection and processing | Pre<sup>wt-fl</sup> | BP<sup>pre</sup> | Pre<sup>A127G</sup> | D1-3 |
| Microscope | Titan Krios |  |  |  |
| Voltage (kV) | 300 |  |  |  |
| GIF Quantum energy filter slit | 20 |  |  |  |
| Detector | Falcon 4i |  | Gatan K3 |  |
| Nominal Magnification | 100,000 × |  | 105,000 × |  |
| Pixel size (Å) | 0.959 |  | 0.83 |  |
| Symmetry imposed | C1 |  |  |  |
| Defocus range (μm) | -1.2– -2.2 |  |  |  |
| Electron exposure (e<sup>-</sup>/Å<sup>2</sup>) | 50 | 50 | 51 | 51 |
| Micrographs (acquired/used) | 8356 |  | 4250 |  |
| Number of extracted particles | 2,335,124 |  | 613,080 |  |
| Number of particles after 3D classifications | 260,038 | 802,830 | 243,014 | 57,917 |
| Map resolution at 0.143 FSC criterion (Å) | 2.75 | 2.52 | 2.84 | 3.12 |
| Local resolution range (Å) | 2.5-4.5 | 2.5-4.5 | 2.5-4.5 | 2.5-4.5 |
| Model refinement |  |  |  |  |
| Atoms | 15891 | 15950 | 15555 | 12684 |
| Residues | 737 | 740 | 720 | 587 |
| CC<sub>mask</sub> | 0.93 | 0.93 | 0.84 | 0.87 |
| Resolution<sub>FSC map vs. model @ 0.5</sub> (Å) | 2.75 | 2.52 | 2.84 | 3.12 |
| R.m.s. deviations |  |  |  |  |
| Bond lengths (Å) | 0.007 | 0.007 | 0.006 | 0.006 |
| Bond angles (°) | 0.904 | 1.003 | 0.762 | 0.861 |
| Validation |  |  |  |  |
| Clash score | 4.13 | 4.19 | 3.91 | 7.82 |
| MolProbity score | 2.35 | 2.35 | 2.33 | 2.58 |

|  |  |  |  |  |  |
| --- | --- | --- | --- | --- | --- |
| Cryo-EM data collection and processing | Pre<sup>G220C</sup> | Pre<sup>G221U</sup> | hyI | hyP1 | hyP2 |
| Microscope | Titan Krios | Titan Actica |  |  |  |
| Voltage (kV) | 300 | 200 |  |  |  |
| Detector | Gatan K3 |  |  |  |  |
| Nominal Magnification | 105,000 × | 100,000 × |  |  |  |
| Pixel size (Å) | 0.83 | 0.82 |  |  |  |
| Symmetry imposed | C1 |  |  |  |  |
| Defocus range (µm) | -1.0– -1.8 |  |  |  |  |
| Electron exposure (e<sup>-</sup>/Å<sup>2</sup>) | 50.15 | 51.21 | 50.12 | 50.46 |  |
| Micrographs (acquired/used) | 5670 | 5308 | 4697 | 6381 |  |
| Number of extracted particles | 1,218,143 | 1,570,760 | 1,125,383 | 848,054 |  |
| Number of particles after 3D classifications | 376,905 | 498,518 | 296,166 | 294,967 | 174,255 |
| Map resolution at 0.143 FSC criterion (Å) | 2.65 | 2.75 | 2.69 | 3.21 | 3.68 |
| Local resolution range (Å) | 2.5-4.5 | 2.5-4.5 | 2.5-4.5 | 3-5 | 2.5-4.5 |
| Model refinement |  |  |  |  |  |
| Atoms | 14510 | 14562 | 15860 | 15635 | 15729 |
| Residues | 673 | 675 | 735 | 726 | 730 |
| CC<sub>mask</sub> | 0.92 | 0.91 | 0.92 | 0.91 | 0.89 |
| Resolution<sub>FSC map vs. model @ 0.5</sub> (Å) | 2.65 | 2.75 | 2.69 | 3.21 | 3.68 |
| R.m.s. deviations |  |  |  |  |  |
| Bond lengths (Å) | 0.007 | 0.006 | 0.008 | 0.008 | 0.008 |
| Bond angles (°) | 0.786 | 0.875 | 1.024 | 1.009 | 1.074 |
| Validation |  |  |  |  |  |
| Clash score | 6.7 | 5.49 | 4.37 | 5.19 | 9.43 |
| MolProbity score | 2.52 | 2.45 | 2.36 | 2.43 | 2.65 |

